# Combined HDAC and eIF4A inhibition: A novel epigenetic therapy for pancreatic adenocarcinoma

**DOI:** 10.1101/2024.06.30.600495

**Authors:** Maryam Safari, Luigi Scotto, Agnes Basseville, Thomas Litman, Haoran Xue, Luba Petrukhin, Ping Zhou, Diana V. Morales, Christopher Damoci, Mingzhao Zhu, Kenneth Hull, Kenneth P. Olive, Tito Fojo, Daniel Romo, Susan E. Bates

## Abstract

Pancreatic ductal adenocarcinoma-(PDAC) needs innovative approaches due to its 12% 5-year survival despite current therapies. We show marked sensitivity of pancreatic cancer cells to the combination of a novel eIF4A inhibitor, des-methyl pateamine A (DMPatA), and a histone deacetylase inhibitor, romidepsin, inducing epigenetic reprogramming as an innovative therapeutic strategy. Exploring the mechanistic activity of this combination showed that with a short duration of romidepsin at low doses, robust acetylation persisted up to 48h with the combination, while histone acetylation rapidly faded with monotherapy. This represents an unexpected mechanism of action against PDAC cells that triggers transcriptional overload, metabolic stress, and augmented DNA damage. Structurally different class I HDAC inhibitors exhibit the same hyperacetylation patterns when co-administered with DMPatA, suggesting a class effect. We show efficacy of this combination regimen against tumor growth in a MIA PaCa-2 xenograft model of PDAC with persistent hyperacetylation confirmed in tumor samples.

**STATEMENT OF SIGNIFICANCE:** Pancreatic ductal adenocarcinoma, a significant clinical challenge, could benefit from the latent potential of epigenetic therapies like HDAC inhibitors-(HDIs), typically limited to hematological malignancies. Our study shows that a synergistic low dose combination of HDIs with an eIF4A-inhibitor in pancreatic cancer models results in marked pre-clinical efficacy, offering a promising new treatment strategy.

## INTRODUCTION

Pancreatic Ductal Adenocarcinoma (PDAC) arises from epithelial cells lining the ducts of the pancreas. PDAC is a highly lethal cancer, ranking third in cancer-related deaths in the United States. Factors contributing to the high mortality rates of patients with PDAC include diagnosis in advanced stages, aggressive tumor biology, and the regular emergence of resistance to existing treatment options, highlighting an urgent need for innovative therapeutic strategies.^1^

The highly proliferative and invasive PDAC epithelial cells rely on frequently mutated drivers and tumor suppressors including KRAS, TP53, CDKN2A, and SMAD4, which disrupt transcriptional regulatory elements and mRNA translation in favor of malignant proliferation.^2–6, 7^ This raises the question of whether targeting factors involved in transcription and translation such as MYC,^8^ histone deacetylases,^9, 10^ or the components of the eukaryotic initiation factor 4F (eIF4F) complex, such as eIF4E or eIF4A could be of value in PDAC.^11, 12^

Histone deacetylases (HDACs) and histone acetyl transferases (HATs) play a crucial role in regulating the levels of histone acetylation that impact gene expression^9^. Dysregulation of acetylation, in part through overexpression of HDACs, has been linked to several cancers including PDAC.^2, 3^ HDAC inhibitors (HDIs) target reversible epigenetic modifications by increasing histone acetylation, chromatin accessibility and transcription of several genes including those that regulate cell growth, apoptosis, or differentiation.^9, 10, 13^ These properties have led to regulatory approvals for HDIs in T-cell lymphomas (CTCL), where persistence of histone acetylation has been shown to be associated with better patient outcomes.^14–17^ However, in solid tumors, HDIs have not yet demonstrated robust anticancer activity.^18^

Tumor cells selectively translate mRNAs crucial for growth, metastasis, and adaptation to the tumor environment.^19, 20^ The majority of such mRNAs are translated via the strictly regulated cap-dependent mechanism that recognizes their highly structured 5′-untranslated regions (UTRs)^21, 22, 23^ with initiation being the rate-limiting step.^22^ During the initiation step, the eukaryotic initiation factor 4F complex (eIF4F) is formed, critical for assembly of the 80s ribosome.^19^ An important subunit of the eIF4F complex is the eukaryotic initiation factor 4A (eIF4A, or DDX2), an RNA helicase essential for unwinding the highly structured 5′ UTRs of cap-dependent mRNAs.^24^ In some cancers including PDAC, alterations in the activity and expression of translation initiation factors such as eIF4A are thought to contribute to tumorigenesis, making these factors attractive therapeutic candidates.^22, 25, 26^ While clinical development of mRNA translation inhibitors has been challenged by toxicities at effective doses, at least two eIF4A inhibitors are currently in clinical trials in various cancer types.^27–30^

Here we report results of combining class I HDAC inhibitors with DMPatA, a novel eIF4A inhibitor, in the treatment of PDAC. DMPatA is a derivative of the natural product pateamine A, which is isolated from the marine sponge *Mycale* sp., and inhibits cap-dependent translation by clamping eIF4A to mRNA.^31–33^ We show this combination is synergistic in killing PDAC cells at low drug concentrations and short treatment periods, the latter selected to emulate the 3.5h half-life of romidepsin. Treatment with this combination remarkably augments HDAC inhibitor activity by enforcing histone hyperacetylation, increasing global transcription and accumulation of co-transcriptional R-loops, replication stress and DNA damage. Our *in vivo* data suggest a safe and effective dose is attainable.

Taken together, this novel epigenetic drug combination has the potential to become a new therapeutic class for pancreatic cancer, enhancing patient outcomes.

## RESULTS

### eiF4A inhibitor DMPatA prevents PDAC cell proliferation and reduces MYC protein levels

Previous studies have reported that inhibitors of translation initiation prevent cell proliferation and reduce levels of critical oncoproteins such as MYC and cyclinD1.^22^ **Figure 1** shows the chemical structure **(1A)**, and crystal structure of DMPatA^33, 34^ bound to the eIF4A/mRNA complex **(1B)**. We first evaluated the effects of DMPatA, a novel inhibitor of eIF4A helicase, on four PDAC cell lines. All cells were sensitive to treatment with IC50s ranging from 2.8 to 10nM **(1C)**. As expected, treatment with both DMPatA^34^ and DMDAPatA,^35^ an earlier analogue of DMPatA, reduced c-MYC levels in all cell lines, with DMPatA achieving comparable effects at lower doses (**1D**). We also evaluated efficacy of another eIF4A inhibitor, zotatifin (eFT226), a rocaglamide derivative that binds to the same binding site as DMPatA on eIF4A,^33, 36^ in MIA PaCa-2 and KLM cells. Our data indicate that DMPatA exhibits significantly greater potency in suppressing proliferation of cells at lower doses compared to zotatifin (**Figure S1A**).

**Figure 1.**
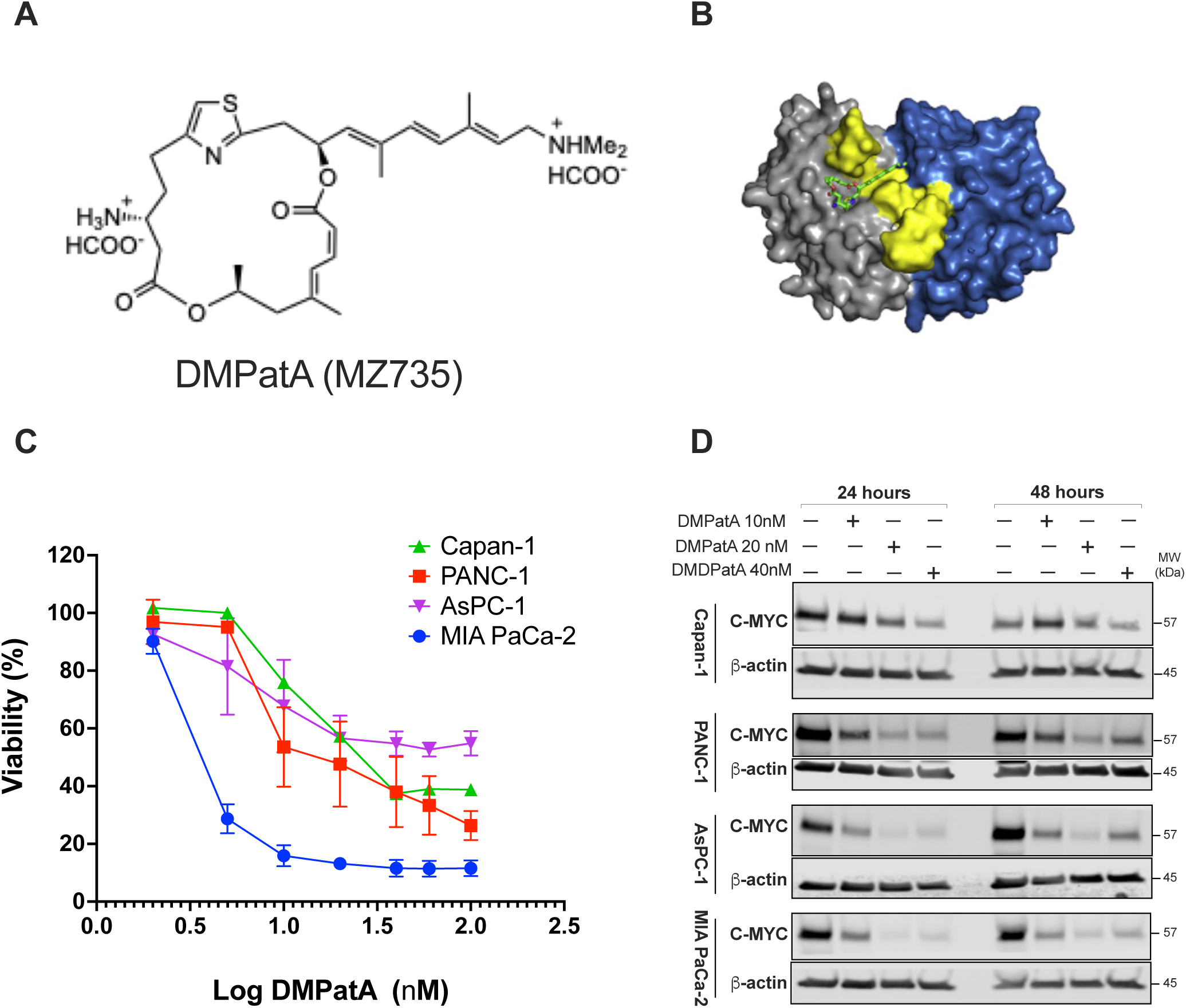
eiF4A inhibitor DMPatA prevents PDAC cell proliferation and reduces MYC protein levels. **A.** Chemical structure of DMPatA (MZ735). **B.** Crystal structure of DMPatA (green) bound to the eIF4A/mRNA complex. The surface of eIF4A1 is shown with the N-terminal domain (gray), C-terminal domain (blue), and RNA (yellow). PDB code: 6XKI **C.** Cytotoxicity graphs of the effect of DMPatA on various PDAC cell lines following 48h of treatment. **D.** Immunoblot of c-MYC protein levels from whole cell extracts of various PDAC cell lines treated with DMPatA and DMDAPatA (Pateamine A). Treatment was for 24- and 48h. Cells were treated with 10 and 20 nM of DMPatA and 40 nM of DMDPatA.

### Combining the HDAC-inhibitor romidepsin with the eIF4A inhibitor, DMPatA, results in synergistic cell death at low drug concentrations

We initially hypothesized that the HDAC inhibitor romidepsin could cause synergistic cell death when combined with DMPatA via a reduction in MYC levels, an effect previously seen separately with both eIF4A inhibitors and with romidepsin in cells harboring mutant KRAS.^37, 38^ We studied the effects of combining DMPatA and romidepsin on cell proliferation in seven PDAC cell lines with different KRAS or BRAF mutations: PANC-1 (*G12D*), PANC 02.13 (*Q61R*), MIA PaCa-2 (*G12C*), Capan-1 (*G12V*), KLM (*G12D*), ASPC1 (*G12D*), and BXPC3 (KRAS WT, BRAF*V487-P492*). To emulate a clinically relevant exposure, romidepsin treatment duration in this and all the experiments described in this manuscript was only 6h ^39–41^, after which it was washed out from combination and single romidepsin samples and the medium replaced with either fresh medium that was drug-free (single agent romidepsin) or contained 2nM DMPatA for an additional 42h (combination). DMPatA control cells were incubated with 2nM DMPatA for 48h. As shown in **Figure 2A**, all the tested cell lines were sensitive to the combination (triangles), albeit to varying degrees, with MIA PaCa-2 cells most sensitive to both DMPatA and combination treatments. Excess over Bliss analysis confirmed synergy of low concentrations of romidepsin with 2nM DMPatA (**Figure 2B).** Additionally, we assessed synergy using a combination of romidepsin with 4 and 10nM zotatifin on MIA PaCa-2 and KLM cells. Similar to the monotherapy results, zotatifin was less potent in combination with romidepsin when compared to 2nM DMPatA (**Figure S1B and C**). Finally, since treatment with DMPatA decreased c-MYC protein levels, we assessed MYC levels after treatment with low doses of both DMPatA (2nM) and romidepsin (15nM). We found reduced c-MYC protein levels following treatment of cells with the combination as well as with DMPatA at lower doses (**Figure 2C)**.

**Figure 2.**
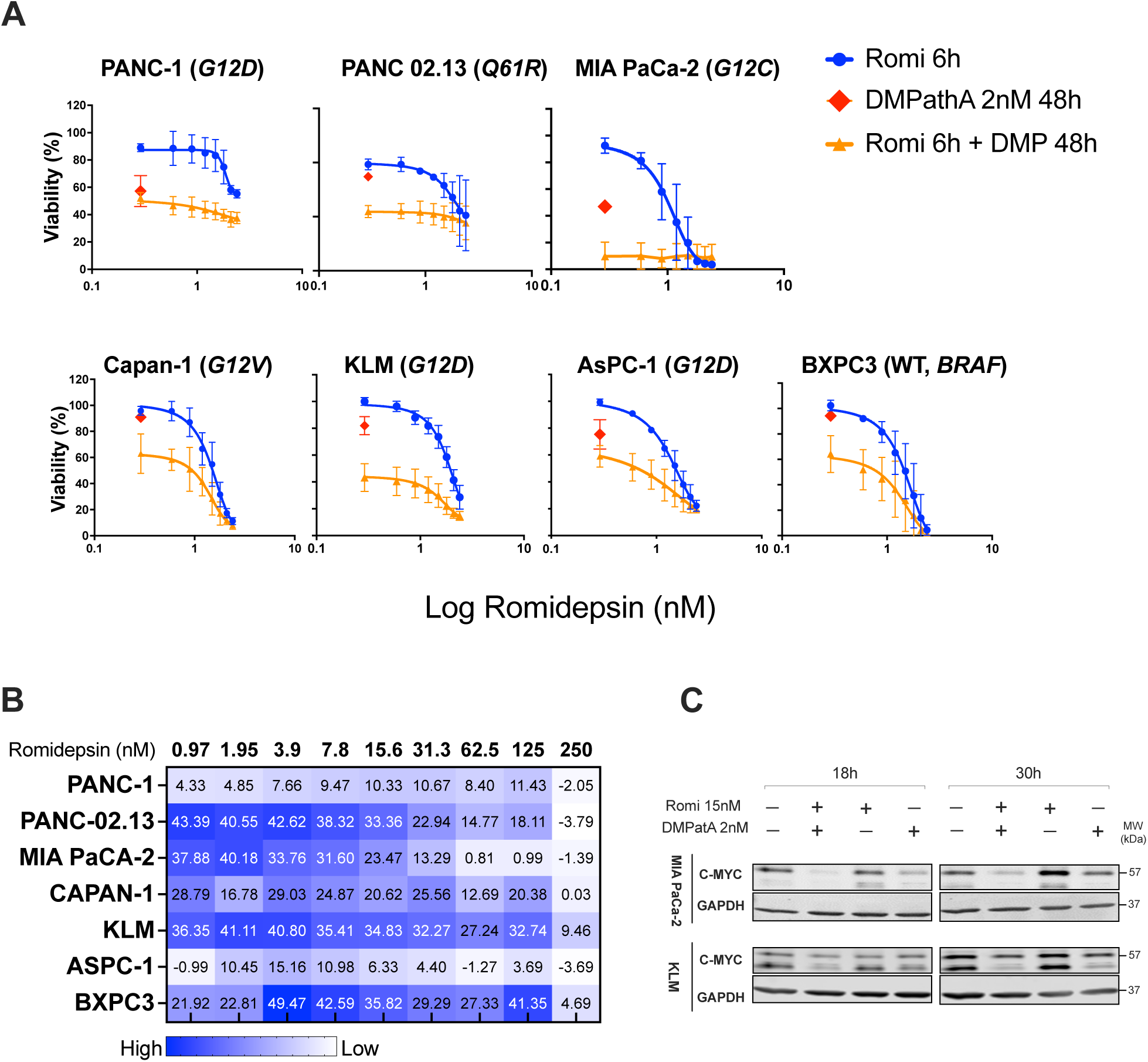
Combining the HDAC-inhibitor romidepsin with the eIF4A inhibitor, DMPatA, results in synergistic cell death at low drug concentrations. **A.** Cytotoxicity graphs of the effect of romidepsin alone at increasing doses (blue), DMPatA alone at 2nM (red diamond), and combination of romidepsin (Romi) with DMPatA at 2nM (orange). Romidepsin treatment was for 6h after which it was washed out and medium replaced with either fresh medium without drug (single agent romidepsin) or with DMPatA (combination) for an additional 42h. DMPatA control cells were incubated with 2nM DMPatA for 48h. **B.** Excess over Bliss synergy (EOB) analysis was used to quantify the degree of synergy when combining various concentrations of romidepsin with 2nM DMPatA. **C**. Immunoblot of c-MYC protein levels from whole cell extracts following treatment. Romidepsin treatment was for 6h (both Romi and Romi+DMPatA) followed by additional incubation without drug (single agent romidepsin) or with DMPatA (combination) for 12 and 24h. DMPatA control cells were incubated with 2nM DMPatA for 18 and 30h.

### Combination treatment impairs metabolism, and MYC overexpression sensitizes cells to the treatment

Because reprograming/augmentation of cellular metabolism by c-MYC is known to support oncogenesis,^42^ we used the Seahorse XF analyzer to assess the impact of the treatments on overall ATP production, glycolysis and mitochondrial oxidative phosphorylation (OXPHOS). We utilized the 6h exposure for romidepsin followed by 18h washout strategy described above. To gain an insight into how different metabolic pathways were affected we first quantified the rate of ATP production from glycolysis and mitochondria simultaneously. Treatment with romidepsin alone and the combination reduced both mitochondrial and glycolytic ATP production (Mito- and Glyco-ATP), while DMPatA treatment mainly impaired the production of ATP by glycolysis (Glyco-ATP) **(Figure 3A)**. This was confirmed by separately assessing glycolysis and mitochondrial OXPHOS pathways by measuring the rate of proton transport across the plasma membrane, proton efflux rate (PER), reflecting glycolysis and the cellular oxygen consumption rates (OCR) reflecting mitochondrial OXPHOS. In MIA PaCa-2 cells, the combination treatment impaired both glycolysis and the mitochondrial OXPHOS and romidepsin impaired mitochondrial function (**Figure 3B and 3C, Left**). However, in KLM cells, which are less sensitive to the treatment within the first 24h, the mitochondrial impairment observed at 18h was less significant (**Figure S2A, Left**). Additionally, to assess the impact of treatment on glutamine oxidation, a primary energy source for cancer cells, especially those driven by c-MYC, we evaluated the ability of glutamine to rescue mitochondrial energy production in cells starved for glucose, glutamine, and pyruvate. In the control cells, glutamine addition increased basal levels of oxygen consumption (ΔOCR), rescuing mitochondrial oxidative phosphorylation by providing glutamine as the energy source. However, treatment with romidepsin alone or the combination and to a lesser extent DMPatA, impaired glutamine rescue of mitochondrial function (**Figure 3C, Right and S2A, Right**).

**Figure 3.**
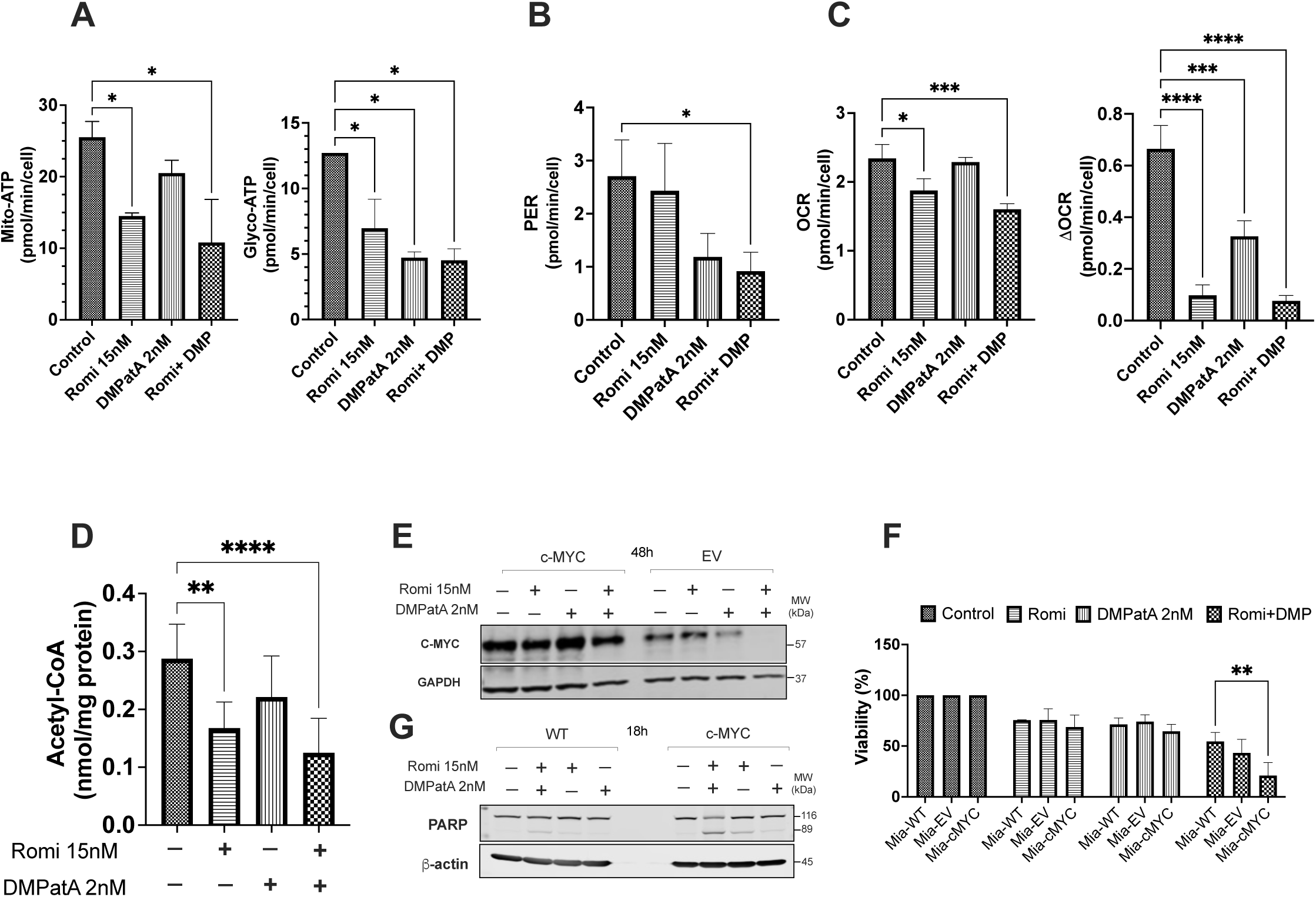
Combination treatment impairs metabolism, and MYC overexpression sensitizes cells to the treatment. The experiments in this figure were conducted in MiaPaCa-2 cells. In Panels **A-D, and G**, cells were incubated for a total of 18h including 6h of 15nM romidepsin (both Romi and Romi+DMP) followed by an additional 12h in drug free medium (Romi) or in 2nM DMPatA (Romi + DMP), or 18 hours in 2nM DMPatA (DMPatA). **A.** Quantification of ATP production rate in MIA PaCa-2 cells. Seahorse XF Real-Time ATP Rate Assay was performed under control conditions or after treatment with either the combination or single agents. **Left** - ATP production rate from mitochondrial respiration (Mito-ATP). **Right** - ATP production rate from glycolysis (Glyco-ATP). At least two biological replicate experiments with n=3 technical replicates were performed**. B.** Impact of treatment on metabolic pathways. Metabolic flux analysis shows basal glycolysis by measuring proton efflux rates (PER), in MIA PaCa-2 cells following treatment with either of the two drugs alone or in combination. At least two independent biological replicate experiments were performed with n=3 technical replicates. **C. Left** - Metabolic flux analysis showing mitochondrial oxygen consumption rates (OCR) reflecting basal mitochondrial oxidative phosphorylation (OXPHOS) in MIA PaCa-2 cells following treatment with single drugs or the combinations. **Right** - Quantification of glutamine consumption. Prior to the metabolic measurements, cells were starved of glucose, glutamine, and pyruvate for 1h after which glutamine was added and response was recorded as ΔOCR (i.e. OCR before and after glutamine addition). **D.** Cellular acetyl-CoA levels were measured in MIA PaCa-2 cells using LC- MS following treatment of cells with single agents or with the combination as above. At least three biological replicate experiments were carried out. **E.** Immunoblot of c-MYC protein levels in whole cell lysate of MIA PaCa-2 cells transfected with a MYC over-expressing (c-MYC) or an empty vector (EV), with or without treatment with single agents or the combination for a total of 48h as in Figure 2. **F.** Cytotoxicity assay of c-MYC and EV MIA PaCa-2 cells following treatment for a total of 24h**. G.** Immunoblot of PARP protein levels in whole cell lysate of wild type and c-MYC over-expressing MIA PaCa-2 cells with or without treatment with single agents or the combination. **Notes:** In all experiments data are expressed as mean ± SD. Statistical significance was determined by one-way ANOVA using Tukey multiple comparisons test in the Prism software. All p-values are relative to control with asterisks as follows: [*], p-value<0.01; [**], p-value<0.002; [***], p-value<0.001; [****], p value<0.0001. Abbreviations: DMP, DMPatA; Romi, romidepsin, PER, Proton Efflux Rate; OCR, oxygen consumption rate.

Previous studies have demonstrated that histone acetylation following treatment with the HDAC inhibitor romidepsin reduces cellular acetyl-CoA levels, a key donor for histone acetylation and an important metabolic intermediate. ^37^ These studies suggested that reduced acetyl-CoA levels might underlie the impaired glutamine utilization. We measured cellular acetyl-CoA levels and observed a significant reduction in cells treated with either romidepsin alone or the combination in both MIA PaCa-2 and KLM cell lines. (**Figure 3D and S2B).**

Finally, given drug treatment especially with the combination, leads to reduced MYC expression, and this could blunt cytotoxicity, we assessed the impact of sustained MYC expression in sensitivity of the MIA PaCa-2 cells to drug treatment. We transfected the cells with an empty and a MYC overexpressing vector (**Figure 3E and S3A**). Maintaining MYC levels with the MYC overexpressing vector led to enhanced sensitivity to the drug combination and further impaired mitochondrial OXPHOS (**Figures 3F** and **3SB**). This finding was confirmed by the increased PARP cleavage observed in c-MYC cells treated with combination, compared to wild-type MIA PaCa-2 cells (**Figure 3G**).

### Combination treatment increased gene expression

A well-known effect of HDAC inhibition is altered gene expression, thought to be due to increased chromatin accessibility following histone acetylation with both gene induction and gene repression having been demonstrated.^37, 43^ We profiled gene expression in MIA PaCa-2 and KLM cells treated with vehicle, each single agent, or the combination for 18h to look for gene expression changes that could provide insights into the treatment effects. Initial analysis showed a significant increase in the number of differentially expressed genes (DEG) in cells treated with the combination compared to the single agents (5773 DEGs for the combination treatment versus 1551 and 1443, respectively, for romidepsin and DMPatA as single agents) (**Figure 4A**). Pathway enrichment analysis associated with the DEGs found *proton motive force-driven mitochondrial ATP synthesis*, *mitochondrial respiratory chain complex assembly*, and *intrinsic apoptotic signaling pathways* significantly enriched following all treatments, in line with the **figures 2 and 3** results documenting cell death and perturbations in metabolism particularly following treatment with the combination. Romidepsin and DMPatA treatment did not target the same genes in these pathways, and the combination induced significantly larger (6 to 8 times) alterations in gene expression than in monotherapy (**Figure 4B, S4A and B, S5A**). Additionally, we noticed enrichment of genes related to *histone modification* in the samples treated with the combination (183 genes) and DMPatA alone (63 genes) suggesting eIF4A could play a role in regulating histone post translational modification **(Figure 4B and S6A**). These genes predominantly encode chromatin modifiers such as histone lysine demethylases (KDMs), histone deacetylases (HDACs), histone acetyltransferases (HATs), and histone methyltransferases (HMTs). Analysis of the chromatin modifier genes revealed a significant overlap in the lysine demethylase (KDM) family when comparing treatment with single agent DMPatA and the combination with increased expression of KDM7A, KDM5B, KDM6B, KDM3A, and KDM4C with both treatments. In addition, HATs (KAT2A and CREBBP), chromatin remodeling factors (ING4 and NCOA3), and HDACs, particularly SIRT1, were among categories of shared genes with altered expression with both the combination and singe agent DMPatA (**Figure S6A**).

**Figure 4.**
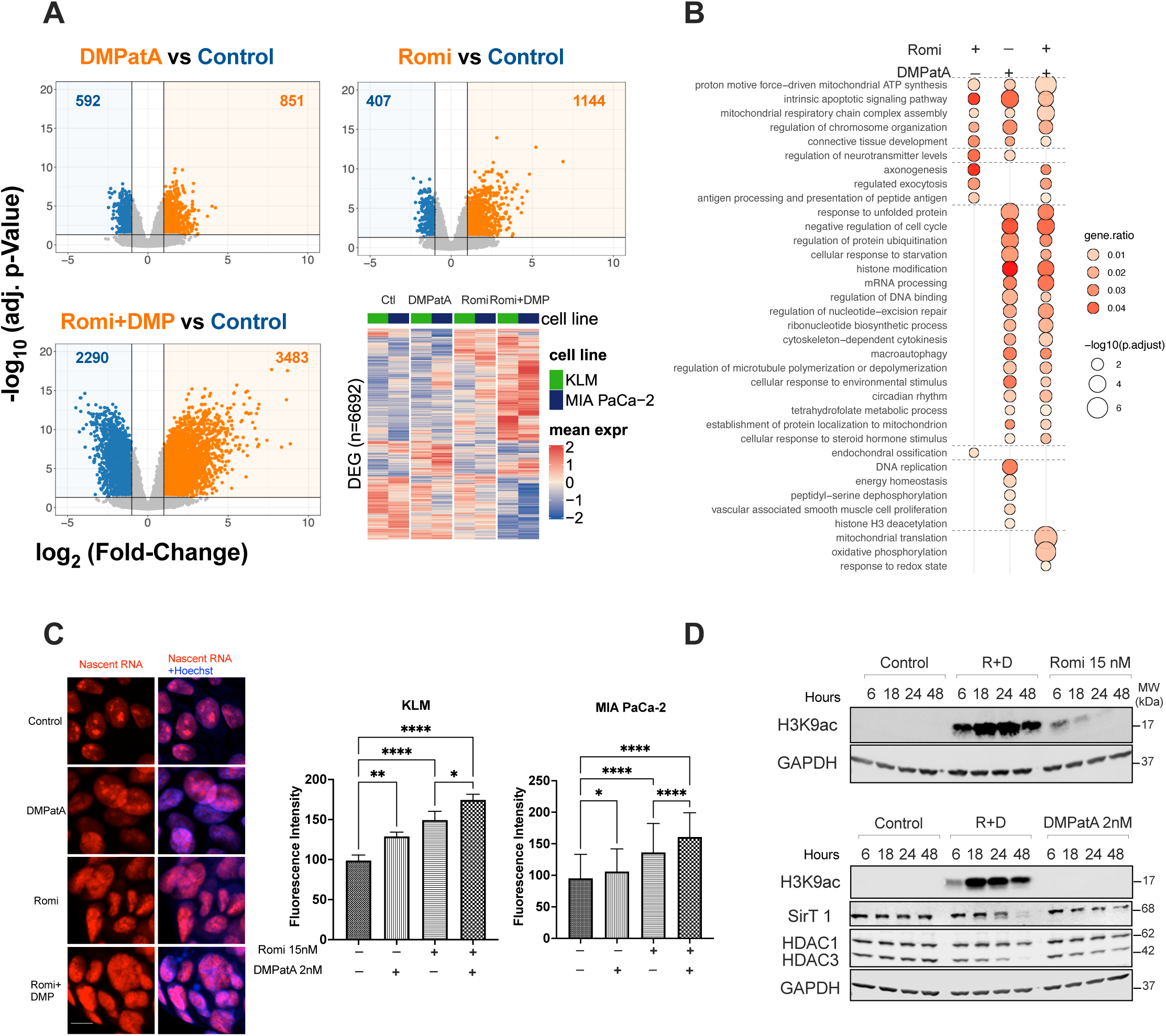
Combination treatment increased gene expression, induced hypertranscription and hyperacetylation. The experiments in Panels **A, B** and **C** were conducted in MiaPaCa-2 and KLM cell lines incubated for a total of 18h, with 6h of 15nM romidepsin (both Romi and Romi+DMP) followed by an additional 12h in drug free medium (Romi) or in 2nM DMPatA (Romi + DMP), or 18h continuously with 2nM DMPatA (DMPatA). **A.** Differentially expressed genes shared by the 2 cell lines after treatment were plotted on Volcano plots (adjusted p-value versus fold change). Variation in expression for the significantly modified genes in a least one condition was plotted on a heatmap (mean triplicate expression for each cell lines). **B.** Pathway enrichment analysis was performed by hypergeometric test, then enriched pathways for the three treatments were filtered with an adjusted p-value <0.05, and GO semantic similarity was used to decrease pathway redundancy and reach a total of 66 pathways from the 429 initial concatenated ones. **C. Left** EU-labeled nascent RNA (red) and Hoechst counterstain (blue) in KLM cells treated with either the single agents or their combination. Scale bar: 1 μm. **Right** Quantification of fluorescence intensities of nascent RNA synthesis in both the KLM and MIA PaCa-2 cell lines. Statistical significance was determined by one-way ANOVA using Tukey multiple comparisons test, [*], p-value<0.01, [**], p-value<0.002, [****], p-value<0.0001. **D.** Immunoblots showing levels of the H3K9ac, SirT1, HDAC1 and HDAC3 in untreated MIA PaCa-2 cells or following treatment with the single agents or their combination for the times indicated. Romidepsin treatment as a single drug or in combination with DMPatA was for 6h. Abbreviations: DMP, DMPartA; EU, 5-ethynyl uridine; Romi, romidepsin.

### Combination treatment induced hypertranscription and hyperacetylation

We quantified transcription and evaluated the global acetylation levels. At low doses (15 nM) and short exposures (6h), single agent romidepsin only modestly increases gene expression, however, the combination of romidepsin and DMPatA markedly increased the number of genes with altered expression levels. We confirmed that the observed increase was due to elevated transcription, as evidenced by the increased output of nascent RNA above normal levels. This was quantified following an 18-hour treatment, (6h for romidepsin), using the incorporation of the modified RNA precursor 5-EU. While an increase in RNA synthesis was seen with romidepsin, and to a lesser extent with DMPatA as single agents, in both KLM and MIA PaCa-2 cells, treatment with the combination significantly increased mRNA synthesis (**Figure 4C**). Since HDAC inhibitors increase histone acetylation and considering the observed increase in gene expression following combination treatment, we assessed the impact of this combination on global histone acetylation. Although DMPatA alone did not impact histone acetylation, its combination with romidepsin led to an unexpected increased H3K9 acetylation both in level and duration that notably exceeded the levels observed with romidepsin alone, again at the low concentration (15nM) and short duration (6h) used in these experiments (**Figure 4D**). Since romidepsin-induced acetylation is primarily due to inhibition of class I HDAC activity, considering the observed histone hyperacetylation with the combination, we examined protein levels of specific class-I HDAC enzymes following treatment with DMPatA alone and in combination with romidepsin. We found a reduction in the levels of HDAC3, SirT1 and phospho-SirT1 proteins with both treatments but notably more evident in combination indicating that at least some of these enzymes rely on eIF4A and cap-dependent translation (**Figure 4D and S6B**). Increased histone acetylation was also observed at other histone marks such as H3K27ac and H2BK9ac modifications (**Figure S7A**). A drug class effect is suggested by the similar pattern of histone hyperacetylation observed with combinations of DMPatA with other HDAC inhibitors, including entinostat and chidamide (**Figure S7B-C**). These results suggested that the main role of DMPatA, at low concentration used in the combination, was to amplify the histone acetylation signal and downstream consequences in inducing cell death, rather than via a separate biological effect.

### Combination treatment induces accumulation of co-transcriptional R-loops

We previously reported that romidepsin-induced histone acetylation leads to R-loop accumulation, increasing genome instability.^44^ R-loops are byproducts of transcription, three strand nucleic acid structures that form during active gene transcription, where the nascent RNA is hybridized to its complementary DNA sequence leaving displaced and vulnerable single-stranded DNA (ssDNA) ^45^. Since treatment of cells with the combination increased global transcription and led to persistent histone hyperacetylation, we postulated the treatment could similarly enhance the accumulation of R-loops and contribute to cell death. Using the S9.6 antibody against DNA–RNA hybrids, we observed colocalization of S9.6 staining with nucleolin after treatment with the combination for 9h (6h for romidepsin). We confirmed the structures as R-loops by their sensitivity to RNaseH1 treatment (**Figure 5A left and right panels**). R-loop accumulation was further confirmed by DRIP-qPCR (**D**NA:**R**NA **I**mmuno**p**recipitation). To select candidate genes for testing R-loop accumulation using DRIP-qPCR, we used the R-Loop DB (http://rloop.bii.a-star.edu.sg) and the R-loopBase (https://rloopbase.nju.edu.cn/index.jsp) databases. We identified genes with increased expression that have R-loop forming sequences (RLFS). After selecting two gene candidates, *VGF and HBA2,* we performed DRIP analysis in MIA PaCa-2 cells. Using the S9.6 antibody to immunoprecipitate genomic DNA with or without RNaseH1 treatment, followed by qPCR, we confirmed a significant increase in the percent of input of these loci in samples treated with romidepsin alone and with the combination (**Figure 5B**). A modest increase in R-loop accumulation was also observed in cells treated with single-agent DMPatA, suggesting the enrichment of genes related to *histone modification* seen in the pathway enrichment analysis and the decrease in HDAC3 and SirT1 protein levels could contribute to increased transcription and R-loop accumulation following eIF4A inhibition. We also examined *SNRNP* as an R-loop negative locus in MIA PaCa-2 cells and found R-loops did not accumulate at this locus (**Figure 5B**). Finally, chromatin immunoprecipitation (ChIP)-qPCR using an antibody against H3K9ac showed that the R-loop–positive loci VGF *and HBA2* were hyperacetylated following treatment with the combination and, to a lesser extent, with romidepsin alone. (**Figure 5C**).

**Figure 5.**
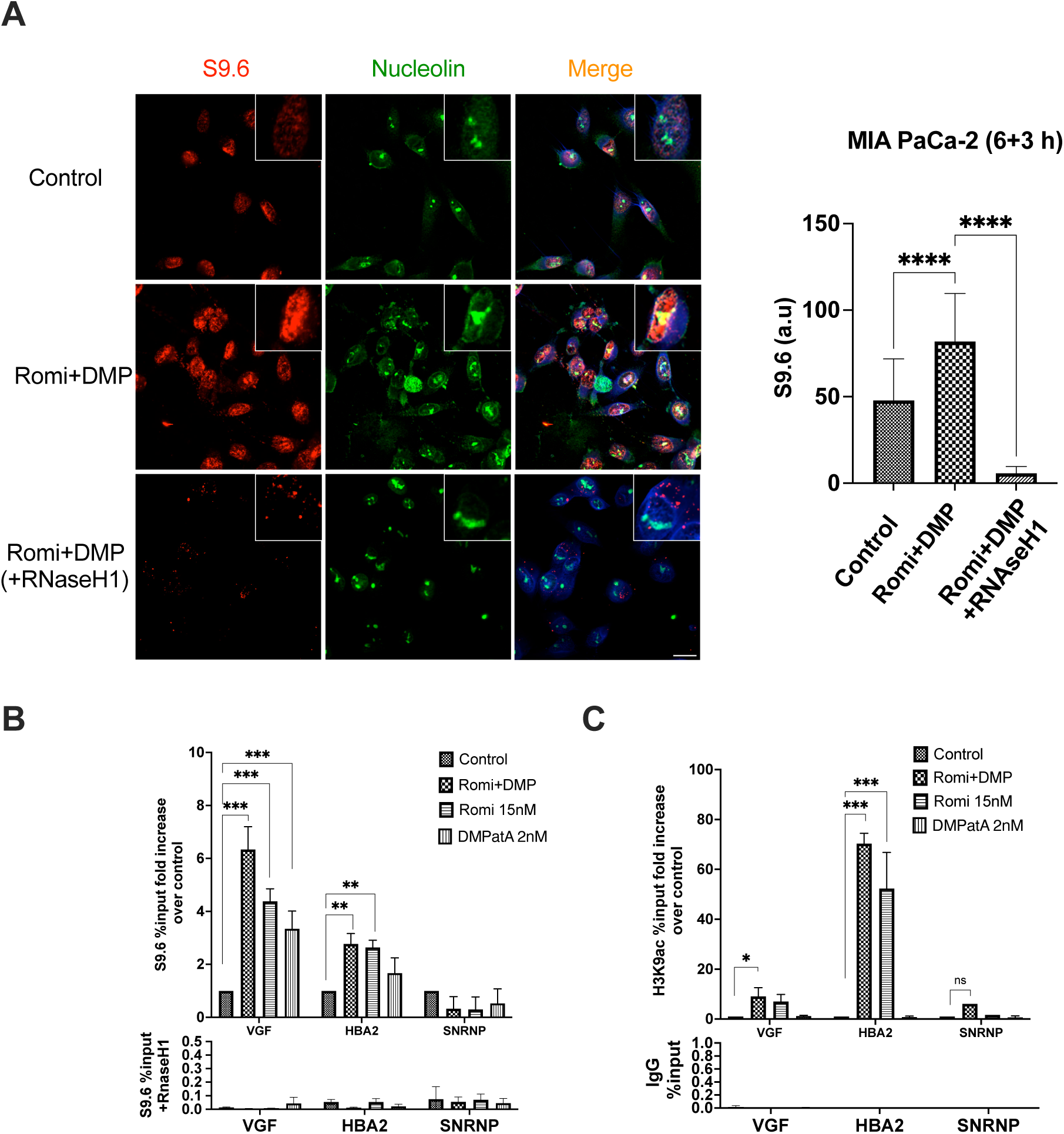
Combination treatment induces accumulation of co-transcriptional R-loops. The experiments were conducted in MiaPaCa-2 cells incubated for a total of 9h including 6h of 15nM romidepsin (both Romi and Romi+DMP) followed by an additional 3h in drug free medium (Romi) or 2nM DMPatA (Romi + DMP) or 9 hours in 2nM DMPatA (DMPatA). **A. Left** Confocal images of DNA–RNA hybrid accumulation in treated cells. Staining was carried out using antibodies against S9.6 (red) and nucleolin (green). Nuclei were stained with DAPI. As seen in the three lower photomicrographs, treatment with RNaseH1 eliminated the DNA–RNA hybrids seen with treatment. Scale bar: 20 μm. **Right** Quantification of fluorescence intensities of nuclear S9.6 immunofluorescence signal in MIA PaCa-2 cells treated with the combination without and with RNaseH1. The S9.6 signal was quantified only in the nuclear regions. The median of the nuclear S9.6 signal intensity per nucleus is shown. Statistical significance was determined by one-way ANOVA using Tukey multiple comparisons test in Prism software, [****], p-value<0.0001. **B.** DRIP–qPCR fold increase in the percent of input values for the *HBA2*, *VGF*, and *SNRPN* loci. Genomic DNA was treated *in vitro* with or without RNaseH1 before immunoprecipitation using the S9.6 antibody. Statistical significance was determined by 3-way ANOVA [**], p-value<0.003, [***], p-value<0.001. **C.** ChIP-qPCR shows fold increase in the percent of input values for the *HBA2*, *VGF*, and *SNRPN* loci. Genomic DNA was immunoprecipitated using H3K9ac antibody. Statistical significance was determined by 3-way ANOVA. [*], p-value<0.03, [***], p-value<0.001. Abbreviations: DAPI, 4’,6-diamidino-2-phenylindole DMP, DMPatA, Romi, romidepsin

### Hyperacetylation and hypertranscription are associated with replication fork stalling and increase in DNA damage

Having observed R-loop accumulation following treatment with the combination, we hypothesized that hypertranscription, persistent hyperacetylation, and R-loop accumulation could increase the incidence of transcription replication conflicts, genome instability, and DNA damage, and contribute to cell death. In response to DNA damage, replication protein A2 (RPA2) is phosphorylated at multiple sites by different protein kinases, including ATM, ATR, and DNA-PK, while the serine threonine-protein kinase, checkpoint kinase 1 (Chk1), is phosphorylated and activated by ATR. We observed accumulation of phospho-RPA2 following treatment with the combination and single-agent romidepsin, and phospho-Chk1 primarily following treatment with the combination (**Figure 6A)**. Looking at replication fork dynamics in both MIA PaCa-2 and KLM cells, we observed significant reduction in the fork speed following treatment after a total of 24h (6h for romidepsin), mainly with the combination and with single-agent romidepsin indicating stalling of replication forks (**Figure 6B).** Consistent with this, in both MIA PaCa-2 and KLM cells, we observed DNA double-stranded damage, indicated by the accumulation of phospho-histone H2AX (γH2AX) protein over time following treatment with the combination, alongside the histone hyperacetylation (**Figure 6C**). This was also confirmed by detection of H2AX foci by confocal microscopy following treatment with single agent romidepsin or the combination (**Figure 6D**).

**Figure 6.**
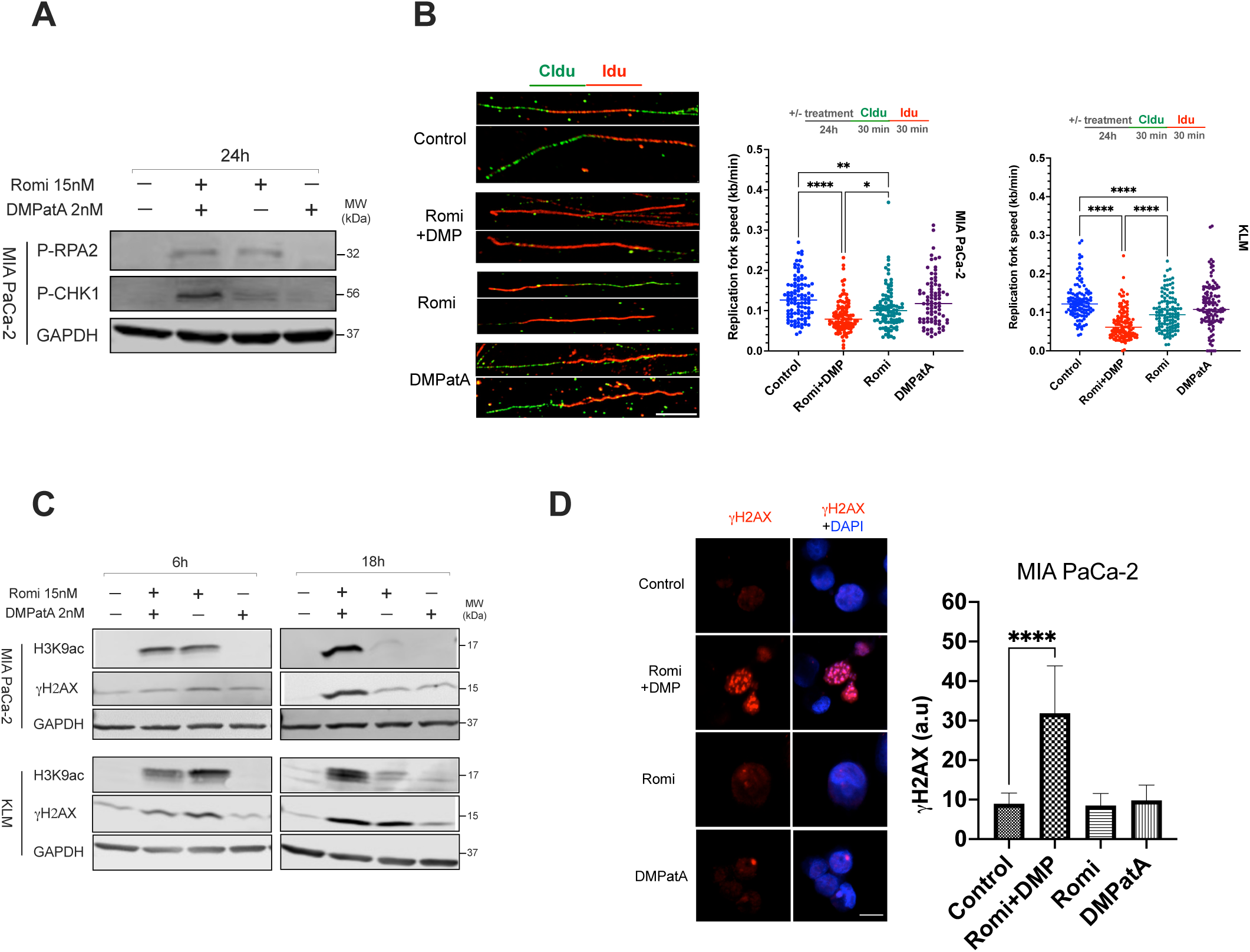
Hyperacetylation and hypertranscription are associated with replication fork stalling and increase in DNA damage. The experiments were conducted in MiaPaCa-2 cells incubated for a total of 24h including 6h of 15nM romidepsin (both Romi and Romi+DMP) followed by an additional 18h in drug free medium (Romi) or 2nM DMPatA (Romi + DMP) or 24h in 2nM DMPatA (DMPatA). **A.** Immunoblot of phosphorylated replication protein A2 (p-RPA2) and phosphorylated checkpoint kinase 1 (p-CHK1) protein levels from whole cell extracts following drug treatment. **B. Left** DNA fiber analysis of untreated control cells or treated cells. Scale bar: 0.5 μm. **Right** Dot plot of IdU to CldU tract length ratios for individual replication forks in the cells treated with the combination or the single agents, The center lines show the medians. To calculate the length in kilobase (Kb) the track lengths were multiplied by a factor 2.59. **C.** Immunoblot of H3K9ac and Ɣ-H2AX protein levels from whole cell extracts following treatment. **D. Left** Confocal imaging of Ɣ-H2AX-labeled (red) and DAPI counterstain (blue) cells. Scale bar: 0.5 μm. **Right** Quantification of nuclear Ɣ-H2AX signal. The median value of at least 50 nuclei signal per experimental condition is indicated. **Note:** Statistical significance was determined by one-way ANOVA using Tukey multiple comparisons test. [*], p-value<0.028, [**], p-value<0.007, [****], p value<0.0001. Abbreviations: DMP, DMPatA; Romi, romidepsin.

### Tolerability and efficacy studies of treatment with the combination in the xenograft model of PDAC

Animal studies were performed following establishment of initial dose levels for the individual drugs. We assessed efficacy of the single agents and the combination in a MIA PaCa-2 xenograft model using a tolerable dose and schedule. We observed regression in tumor volume with single agent romidepsin or DMPatA and larger reductions with the combination (**Figure 7A and B**).

**Figure 7.**
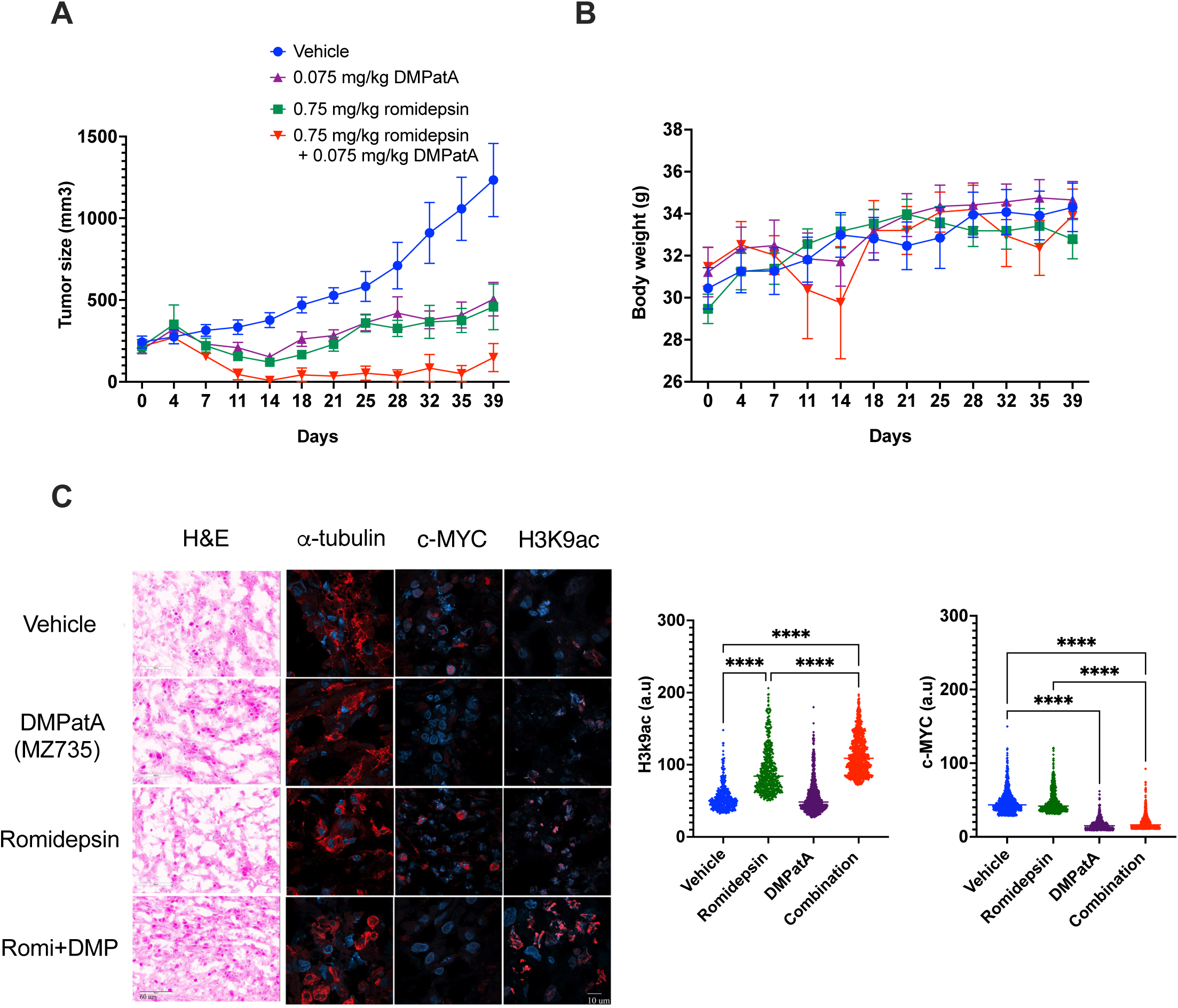
Tolerability and efficacy studies of treatment with the combination in the xenograft model of PDAC. MIA PaCa-2 xenograft model of PDAC in male athymic nude mice. **A.** Tumor growth in mice treated with the indicated concentrations of combination, romidepsin and DMPatA using IP route and vehicles described in the materials and methods section. **B.** Weight of mice treated with the combination and single agents following treatment. **C.** Frozen sections from MIA PaCa-2 xenograft tumor tissues were subjected to either H&E staining or immunohistochemical analysis using antibodies against H3K9ac, c-MYC, and α-tubulin. H&E-stained images are captured at 40x magnification. Scale bar: 60 μm. Fluorescence images were captured at 63x magnification using confocal microscopy. Scale bar: 10 μm. The fluorescent signal from H3K9ac and c-MYC antibodies were quantified. To quantify fluorescence intensity at least 500 cells per condition were measured for each antibody at 40x magnification captured images (see Figure S8). The median of the signal intensity is shown. DAPI staining shows nuclei of cells. Statistical significance was determined by one-way ANOVA using Tukey multiple comparisons test in the Prism software**, [**\*\*\*\*], p value<0.0001. Abbreviations: DMP, DMPatA; Romi, romidepsin.

This suggests that the remarkable synergy observed *in vitro* can translate into an *in vivo* model. To determine whether our *in vitro* mechanistic observations would be observed in tumors, we collected snap-frozen tumor samples from mice treated with vehicle, single agents, and the combination at the end of the study, 72h after the last dosing. These samples were sectioned and the levels of H3K9ac, c-MYC, and α-tubulin proteins in tumor tissues were assessed by immunohistochemistry. In line with our *in vitro* observation of increased histone acetylation in cells treated with the combination, levels of H3K9ac were significantly higher in tumors from mice treated with the combination compared to those that received romidepsin monotherapy. Similarly, we observed lower levels of c-MYC protein expression in the tumor samples from mice treated with DMPatA and the combination, in line with our *in vitro* observation that following treatment with DMPatA and the combination expression of c-MYC protein decreased in PDAC cell lines. No difference in α-tubulin expression, used as an internal control, was observed in mice tumor tissues across vehicle and treatment groups. (**Figure 7C, and S8A**).

## DISCUSSION

Pancreatic ductal adenocarcinoma (PDAC) is a highly lethal solid tumor that readily develops resistance to standard therapies and needs innovative treatments. While new drugs are in development to target the common KRAS mutations, they are not curative, and drug resistance appears to emerge in patients. ^46^ Epigenetic changes along with core driver mutations in KRAS, p53, and other genes play roles in PDAC tumor evolution and metastasis.^2–4^ Epigenetic therapies, particularly HDIs, could offer a promising approach for PDAC treatment by targeting reversible epigenetic modifications that drive tumor heterogeneity and therapy resistance.

Herein, we report results of a novel combination comprising romidepsin, a class I HDAC inhibitor and DMPatA (MZ735), an eIF4A inhibitor. Treatment with this combination increased metabolic stress and induced cell death in PDAC cell lines with diverse KRAS/BRAF mutations, while also prolonging histone acetylation compared to a low dose, short exposure to romidepsin alone. Histone hyperacetylation caused hypertranscription, accumulation of co-transcriptional R-loops, induction of replication stress and double-strand DNA damage in PDAC cells. Using a MIA PaCa-2 xenograft model, our *in vivo* experiments showed marked suppression of tumor growth upon treatment with the combination, suggesting this combination has potential as a first epigenetic therapy for pancreatic cancer.

HDAC inhibitors, initially discovered for their anti-tumor ability by promoting tumor cell differentiation, ^47, 48^ were later found to target HDAC enzymes, causing acetylation of histones and non-histone proteins.^49, 50^ Multiple studies have shown that HDAC inhibition exploits the vulnerabilities of tumor cells promoting tumor-specific cell death attributed to activation of genes regulating differentiation, apoptosis, generation of ROS, and DNA damage. ^37, 44, 51–54^ Additionally, they promote cell cycle arrest by inducing acetylation at the promoter of genes such as CDKN1A, leading to increased p21^WAF1/CIP1^ expression.^51, 55, 56, 57, 58, 59^ During cell cycle, maintaining precise histone modification patterns is essential for maintaining mitotic fidelity.^60, 61^ KRAS-mutant cells are particularly dependent on mitotic fidelity; disrupting mitosis in these cells leads to cell death. ^62^ Disturbing these patterns by HDAC inhibitor-induced acetylation could explain in part the enhanced sensitivity of KRAS mutant cells to HDIs.^37^ These findings underscore the potential clinical relevance of HDAC inhibitors but also highlight their challenges in treating solid tumors and argue for combination therapies to enhance their effectiveness.

Studies on eIF4A inhibitors have shown that a subset of proteins involved in cell cycle regulation, proliferation, and apoptosis, including MYC, Cyclin D1, and BCL-2, rely on cap-dependent protein translation and its inhibition induces apoptosis.^21, 22^ Although various tumors types are sensitive to eIF4A inhibition, toxicities have hindered development of these agents. This encouraged us to develop combination regimens that would use lower doses.^63^

Our studies validated in PDAC cells previous findings on the effects of eIF4A and HDAC inhibitors as single agents, including decreased c-MYC translation. We hypothesized that combining the known downregulation of c-MYC at protein level, via translation, and acetyl-CoA levels, by HDAC inhibition would induce metabolic stress based on the key roles of c-MYC and acetyl-CoA in cellular metabolism. Non-toxic concentrations of both agents in combination caused metabolic stress and cell death and demonstrated an unexpected magnitude of synergy in PDAC cell lines. When we investigated the mechanism underlying the remarkable synergistic effect, we found that the eIF4A inhibitor – HDI combination potentiated the epigenetic effects, i.e., persistent histone hyperacetylation and its ensuing downstream effects.

Upon examining the gene expression profile, we observed an increase in global gene expression with the combination treatment, confirmed by elevated levels of nascent RNA, indicating hypertranscription. Hypertranscription, a concept from stem cell biology, refers to an increase in nascent RNA production beyond normal cellular levels.^64, 65^ Oncogene-induced hypertranscription is linked to R-loop accumulation, heightened transcription-replication conflicts, replication stress, and DNA damage.^66^ Hence, targeting oncogene-induced replication stress has emerged as a promising therapeutic approach.^64^ HDAC enzymes play a role in maintaining replication fork progression.^67^ Inhibition or loss of HDAC1/2 and 3 has been shown to slow the replication fork speed and increase replication stress.^67, 68^ While the role of HDACs in maintaining replication fork stress is established, the impact of histone hyperacetylation-induced hypertranscription in inducing replication stress has not been thoroughly investigated. Our data demonstrate that the combination treatment induces hypertranscription, leading to replication stress, supporting this phenomenon. Importantly, this combination results in a higher degree of hyperacetylation, replication stress, and double-stranded DNA damage compared to the single agent HDAC inhibitor, romidepsin. Furthermore, our data highlight the involvement of both HDACs and eIF4A/cap-translation in preventing transcription-replication conflicts, for which a better understanding can have important therapeutic implications.

RNA-seq analysis of cells treated with the combination or with single-agent DMPatA found mitochondrial respiration and apoptotic signaling related genes among the most impacted. Additionally, genes directly involved in pathways that regulate histone modifications are among the significantly enriched pathways in cells treated with the combination and DMPatA alone. This suggests that the translation of these regulatory proteins may be cap-dependent and provides insights into how DMPatA may contribute to sustaining the hyperacetylation initiated by HDAC inhibitors.

We show that MYC overexpression further sensitizes cells to this combination. MYC activation occurs in 43% of advanced PDAC cases and is associated with a more aggressive tumor behavior.^69, 70^ In cancer cells with elevated c-MYC levels, c-MYC enhances transcription across the entire genome, not just MYC target genes. It does so by accumulating in the promoters of active genes and increasing transcription.^71, 72^ It is therefore likely that MYC activated cells exhibit elevated levels of inherent transcriptional stress, potentially causing further vulnerability to the combination treatment following histone hyperacetylation. Consistent with this we observed increased levels of DNA-damage in c-MYC-overexpressing MIA PaCa-2 cells following combination treatment (Figure S8B). Additionally, a recent study revealed that cells with MYC activation exhibit a heightened need for lipid synthesis to sustain increased proliferation, resulting in a critical dependence on expression of mitochondrial citrate production and transport genes, which generate the acetyl-CoA needed for lipid synthesis. ^73^ Given this, the increased sensitivity of MYC-overexpressing cells could be explained in part by the significant reduction in acetyl-CoA levels caused by combination treatment in cells dependent on lipid synthesis. Interrogating MYC expression as a biomarker for this combination in the clinic could address these possibilities.

We also investigated the effects of combining a different eIF4A inhibitor, zotatifin, with romidepsin, as well as combining other class 1 HDAC inhibitors, such as entinostat and chidamide, with DMPatA. In the former, we observed that zotatifin demonstrated reduced potency, compared to DMPatA, in PDAC cells both alone and in combination. However, the latter indicated that a potential common HDAC regulatory mechanism is targeted when these HDAC inhibitors are combined with DMPatA.

Lastly, our *in vivo* studies have shown that safe and effective doses of this combination can be administered in a murine model, indicating its potential for further development and clinical application.

In conclusion, our results suggest that combining romidepsin and other HDIs with the eIF4A inhibitor DMPatA may offer a promising strategy as a novel epigenetic therapy for pancreatic cancer. Our studies suggest that c-MYC may serve as an important biomarker for this therapy. This combination shows significant antitumor activity in both *in vitro* and *in vivo* models of pancreatic cancer. Further studies are needed to elucidate the precise mechanism of action of this combination and will further enable translation of these findings into the clinic.

## MATERIALS AND METHODS

### Reagents and Cell Culture

Romidepsin (depsipeptide, NSC 630176) was obtained from (Sigma-Aldrich # SML1175) and DMPatA (MZ735) was provided by Dr. Daniel Romo, Baylor University. Entinostat (#S1053) and Chidamide (#S8567) were purchased from Selleckchem. Zotatifin (eFT226, # HY-112163) was obtained from MedChemExpress.

MIA PaCa-2, KLM, ASPC1, PANC-1, PANC 02.13, and Capan-1 cell lines were purchased from American Type Culture Collection. BXPC3 was a kind gift from Dr. Kenneth Olive, Columbia University. All cell lines were regularly authenticated by Genetic Resources Core Facility (GRCF) at Johns Hopkins University. Cells were cultured in appropriate cell culture medium supplemented with 10% fetal bovine serum, 100-units/L penicillin-streptomycin and 1% glutamine. Additionally, Miapaca-2 cells required 2.5% horse serum (Gibco Laboratories, Grand Island, USA).

In all cases, regardless of the duration of the experiment, in both the combination and HDI alone, cells were exposed to romidepsin or other HDI for 6 hours, followed by fresh media without romidepsin. Cells were exposed to the eIF4A inhibitor for the entire duration of the experiment, during the initial 6h, and during the remaining time in romidepsin-free media.

c-MYC-expressing MIA PaCa-2 cells were generated by transfecting wild type MIA PaCa-2 cells with 2μg of pCDNA3mycIRESGFP DNA using Lipofectamine 2000 transfection reagent (Invitrogen, Cat # 11668027). The pCDNA3mycIRESGFP plasmid was created by introducing the BamHI-XhoI fragment of pWZL Blast myc (Addgene plasmid #10674) containing the c-Myc cDNA in BamHI-XhoI restricted pcDNA3.1(+) IRES GFP (Life Technologies, Carlsbad, USA).

### Cell Viability Assay

Cell viability was determined using CellTiter-Glo® reagent (Promega, # G7570) following the manufacturer’s instructions.

### RNA-seq gene expression analysis

Total RNA was extracted using RNeasy mini kit (Qiagen # 74104) from cells harvested after 18h of treatment. RNA library preparation and sequencing was performed at Columbia University’s Sulzberger Genome Center. Briefly, poly-A pull-down was used to enrich mRNAs from total RNA samples, followed by library construction using Illumina TruSeq chemistry. Libraries were then sequenced using Illumina NovaSeq 6000. Abundance of transcripts was estimated by pseudoalignment using kallisto, and downstream analysis was performed using R (v4.3.1) and the indicated packages. Count matrix was normalized using DESeq2 (v1.40.2), followed by a variance stabilizing transformation. Low expression genes (inferior to 5 in more than 85% samples) were filtered out. Differentially expressed genes were determined using linear regression method (limma v3.56.2), and genes with |log2FC| ≥1 and adjusted p-value <0.05 were plotted accordingly (ggplot2 v3.5.0 and ComplexHeatmap v2.16.0). Pathway enrichment was estimated by hypergeometric test (ClusterProfiler v4.8.3) using gene ontology biological process (GO BP) database, with all RNAseq detected genes as gene universe. Pathways with a Benjamini–Hochberg adjusted p-value <0.05 for each treatment were concatenated together, then redundancy in filtered pathways was identified by semantic similarity (Wang distance method) using rrvgo (v1.12.2), and pathway number was reduced accordingly to reach a total of 50 pathways by increasing similarity threshold (from 0.7 to 0.95 in increments of 0.1). Genes associated to some selected pathways before redundancy-based reduction were represented on chord diagram (circlize v0.4.16). The RNA-seq data have been deposited in NCBI’s Gene Expression Omnibus (GEO).

### Labeling of nascent RNA by EU

To detect changes in the global RNA transcription we used Click-iT RNA Alexa Fluor 594 Imaging Kit (Invitrogen, #C10330). Briefly, cells were cultured in complete media and pulsed for 60 min with 5-ethynyl uridine (EU) at a final concentration of 1 mM. After fixation and permeabilization, the 1x Click-iT reaction cocktail was applied for 30 minutes at room temperature. DNA was stained with Hoechst 33342. Images were captured at 40X magnification using a Zeiss Confocal Microscope (Zeiss LSM 700), and Fiji software (ImageJ, version: 1.54f, NIH, USA) was used for image analysis.

### Western blotting

Western blotting was carried out as previously described^39^. Membranes were probed with primary antibodies and secondary antibodies (See table S2). Signal quantified using the Odyssey CLx imaging system (all from LI-COR Biosciences).

### DNA fiber assay

DNA fiber assays were performed using a modified version of protocols described in^74, 75^. Briefly MIA PaCa-2 and KLM cells were seeded at 0.3×10^6^ cells. following treatment cells were pulse-labeled with 25μM CldU (Sigma-Aldrich, USA) for 30 minutes at 37°C with 5% CO_2_, followed by PBS washing. Subsequently, cells were pulse-labeled with 250μM IdU for an additional 30 minutes. After labeling, cells were counted using a cell counter (Nexcelom Bioscience, USA). Two thousand cells were spotted on a microscope slide (Fisher Scientific, USA) and lysed. Slides were then fixed, DNA denatured, and blocked in 5% BSA for 40 min. DNA fibers were immunolabeled using mouse and rat anti-BrdU antibodies and incubated overnight at 4°C. The slides were subsequently incubated with secondary antibodies, goat anti-mouse Cy3 conjugate and goat anti-rat Alexa Fluor 488, for 1h at room temperature. For antibodies, see table S2. After washing, coverslips were mounted, and images were captured at 60X magnification using a Zeiss Confocal Microscope (Zeiss LSM 980). Image analysis was performed using Fiji software (ImageJ, version: 1.54f, NIH, USA).

### Immunocytochemistry of R-loops

R-loop staining was performed as previously described^76^. Cells were incubated with primary antibodies, anti-DNA-RNA hybrid [S9.6] antibody and anti-nucleolin antibody (table S2). Subsequently, secondary antibodies, Alexa Fluor 488 goat anti-mouse or Alexa Fluor 594 goat anti-rabbit used. For antibodies, see table S2. DNA was counterstained with DAPI (Thermo Scientific #62248). Images were acquired at 40X magnification using a Zeiss Confocal Microscope (Zeiss LSM 700), and Fiji (ImageJ) software (version: 1.54f, NIH, USA) was employed for image analysis.

### DNA–RNA immunoprecipitation (DRIP)

DRIP assay was performed according to the method described earlier^77^. Briefly, following gentle extraction of the genomic DNA, cells were subjected to digestion using a cocktail of restriction enzymes, treated with or without RNaseH (New England Biolabs # M0297L). DNA-RNA hybrids were then immunoprecipitated from the digested genomic DNA using S9.6 antibody. Quantitative PCR was performed at the R-loop positive loci *VGF, and HBA2* genes and R-loop negative locus SNRPN gene with the primers listed in table S1.

### ChIP-qPCR

The chromatin immunoprecipitation (ChIP) assay was performed using the Zymo-Spin ChIP kit (Zymo Research, CA #D5210) according to manufacturer’s instructions. After treatment, cells were harvested, and chromatin was cross-linked with formaldehyde, then mechanically sheared by sonication on ice. Protein-DNA complexes were precipitated using control immunoglobulin G (normal Rabbit IgG) and anti-Acetyl-Histone H3 (table S2). Subsequently, ChIP-DNA was eluted, reverse-cross-linked, and purified. Quantitative PCR was performed in technical triplicates using SsoAdvanced Universal SYBR Green Supermix (Bio-Rad, # 1725272) and specific primers for, *VGF, HBA2* and SNRPN genes (see table S1 for primer details).

### Immunofluorescence staining-DNA damage detection

For detecting DNA double-strand damage, 0.3×10^6^ cells were seeded on coated coverslips (Neuvitro, # GG-22-PDL) and treated as required. Cells were then washed, permeabilized with cold methanol, and blocked with 5% BSA. Anti-phospho-Histone H2A.X (Ser139) antibody used for staining. After overnight incubation cells were washed, and Alexa Fluor 488 goat anti-mouse secondary antibody was added for 1h. Coverslips were mounted on glass slides and DAPI (Thermo Scientific #62248) was used to stain DNA. Images were obtained at 40X magnification with a Zeiss Confocal Microscope (Zeiss LSM 700), and Fiji software (ImageJ, version: 1.54f, NIH, USA) was used for image analysis.

### Metabolic flux analysis (Agilent Seahorse XF assay)

We used XF96 Extracellular Flux analyzer (Seahorse Bioscience) to measure ATP production rate (Cat# 103591-100, Agilent), glycolysis (Cat# 103020-100) and mitochondrial OXPHOS (Cat# 103015-100). Data were normalized by cell number using the CyQUANT Cell Proliferation Kit (Thermo Fisher Scientific). For all measurements, 20,000 cells were plated in XF96 cell culture microplates and treated with combination or single agents for a total of 18h. Afterward, growth media was replaced with bicarbonate-free assay media (Seahorse Biosciences) and incubated at 37°C for 1h in a CO2-free incubator. **O**xygen **c**onsumption **r**ate (OCR) reflecting mitochondrial OXPHOS was measured under basal conditions or following the addition of Oligomycin (1μM), FCCP (0.5μM), and Rotenone/Antimycin A (0.5μM), following the manufacturer’s protocol. **P**roton **e**fflux **r**ate (PER), reflecting glycolysis, was measured under basal conditions and following the addition of glucose (10mM), Oligomycin (1μM), and the glucose analog 2-deoxyglucose, 2DG (50mM), following the manufacturer’s protocol.

### Quantification of cellular Acetyl-CoA levels

Cells were treated with combination or single agents for a total of 18h. Following treatment cells were harvested on ice, washed, and scraped out under liquid nitrogen and aliquot of the cell pellet was collected and subjected to protein quantification.

For Acetyl-CoA extraction, a 100 µL aliquot of the thawed pellet was spiked with 13C2-acetyl-CoA as an internal standard, extracted with 400 µL of methanol, vortexed, and centrifuged at 18.0xG for 5 minutes at 4°C. The methanol supernatant (400 µL) was then dried under nitrogen at 35°C and reconstituted in 100 µL of 10mM ammonium bicarbonate (pH 9.5) for LC/MS analysis. Calibration curve standards were similarly prepared as the cell pellets. Acetyl-CoA quantification by LC/MS was performed on an Agilent 1290 Infinity UHPLC/6495 triple quadrupole mass spectrometer, using multiple reaction monitoring. Data processing employed Mass Hunter software (Agilent), and calibration curves (R2 ≥ 0.99) were fitted using either linear or quadratic models with 1/X or 1/X^2 weighting.

### *In vivo* experiment

Animal studies were performed in accordance with procedures approved by the Institutional Animal Care and Use Committee (IACUC) protocol number AC-AABD2601, at the Columbia University Oncology Precision Therapeutics and Imaging Core (OPTIC). Male athymic nude mice were obtained from Jackson Labs. Mice were implanted with 2.5x10^6^ MIA PaCa-2 cells, injected subcutaneously into the right flank with 50% Matrigel. Once tumors reached 150-250 mm^3^ in volume, mice were randomized and enrolled into the study. Each group had a total of eight mice per treatment group. Treatment groups included: Vehicle, Romidepsin 0.75 mg/kg, DMPatA (MZ735) 0.075 mg/kg, Combination: Romidepsin 0.75 mg/kg + DMPatA 0.075 mg/kg. Mice were dosed once every four days intraperitoneally (IP), for 2 weeks initially followed by once every seven days dosing for an additional 5-6 weeks. Formulations for DMPatA were prepared utilizing a solution comprising 10% Ethanol, 10% Cremophor EL, and 80% 4% glucose in PBS. Initially, DMPatA was dissolved in ethanol, followed by the sequential addition of the remaining constituents of the vehicle formulation. Formulations for romidepsin were made using 5% Ethanol and 5% Propylene Glycol, and 90% USP Saline.

### Immunohistochemistry of tumor tissues

Frozen tumor tissues from the *in vivo* experiment were sectioned and stained at the Columbia University Molecular Pathology Shared Resource. Briefly, sections underwent three PBS washes and were blocked using 10% normal goat serum in PBS for 40 min. Primary antibodies Histone H3 (acetyl K9), C-MYC, and alpha tubulin, were applied and incubated for 2h at room temperature. After additional washes, sections were incubated with secondary antibodies Donkey anti-rabbit Alexa 594 or Donkey anti-mouse Alexa 594, for 1h at room temperature. Sections were mounted with DAPI medium. Slide Images were captured at 60X magnification using a Zeiss Confocal Microscope (Zeiss LSM 980). Image analysis was performed using Fiji software (ImageJ, version: 1.54f, open-source image processing software, NIH, USA).

## ACKNOWLEDGMENTS

This research was funded by the Translational Research Grant from Pancreatic Cancer Action Network (PanCAN) awarded to Susan E. Bates (20-65-BATE).

Authors would like to acknowledge the James J. Peters Bronx VA Medical Center for providing laboratory space and support, as well as the efforts of the Bronx Veterans Medical Research Foundation.

This research was funded in part through the NIH/NCI Cancer Center Support Grant P30CA013696 and used the Genomics and High Throughput Screening Shared Resource, Molecular Pathology Shared Resource, and Oncology Precision Therapeutics and Imaging Core (OPTIC) of the Herbert Irving Comprehensive Cancer Center at Columbia University.

Metabolomics studies for Acetyl-CoA quantification were performed by the Metabolomics Core (RRID:SCR_022381) in the Penn Cardiovascular Institute, supported in part by NCI P30 CA016520.

Partial support by NIGMS to D.R. (R35 GM134910) is gratefully acknowledged.

Authors would like to acknowledge the support of the Bioinformatics Core Facility BiRD, part of Biogenouest and the Institut Français de Bioinformatique (IFB; ANR-11-INBS-0013), for providing access to their resources and invaluable technical assistance.

Authors would like to thank Adam Abramowitz for his invaluable assistance in identifying gene candidates for the DRIP-qPCR experiment, leveraging his expertise in data analysis.

