## Supplemental figure legend-figures and tables for "Combined HDAC and eIF4A inhibition: A novel epigenetic therapy for pancreatic adenocarcinoma"

#### Supplementary figure legends:

##### Figure S1.

**A.** Cytotoxicity graphs of the effect of various concentrations of zotatifin (eFT226) on MIA PaCa-2 and KLM cell lines following 48h of treatment. **B.** Cytotoxicity graphs of the effect of romidepsin alone at increasing doses (blue), zotatifin alone at 4nM (red diamond), and combination of romidepsin with zotatifin at 4nM (orange). Romidepsin treatment was for 6h after which it was washed out and medium replaced with fresh medium without (single agent romidesin) or with zotatifin for an additional 42h (combination). **C.** Cytotoxicity graph of the effect of romidepsin alone at increasing doses, zotatifin alone at 10nM and combination of romidepsin with zotatifin at 10nM on MIA PaCa-2 cells. Treatment was as mentioned in B.

##### Figure S2.

The experiments in this figure were conducted in KLM cells. Cells were incubated for a total of 18h including 6h of 15nM romidepsin (both Romi and Romi+DMP) followed by an additional 12h in drug free medium (Romi) or 2nM DMPatA (Romi + DMP) or 18 hours in 2nM DMPatA (DMPatA). **A. Left**- Metabolic flux analysis showing mitochondrial oxygen consumption rates (OCR) reflecting basal mitochondrial oxidative phosphorylation (OXPHOS) in KLM cells following treatment with single drugs or the combinations. **Right** - Quantification of glutamine consumption. Prior to the metabolic measurements, cells were starved of glucose, glutamine, and pyruvate for 1h after which glutamine was added and response was recorded as ΔOCR (i.e. OCR before and after glutamine addition). Statistical significance was determined by one-way ANOVA, [***] p value<0.0001, [*] p value<0.02. **B.** Cellular acetyl-CoA levels measured in KLM cells following treatment of cells with the combination or with single agents using LC-MS. Statistical significance was determined by two-way ANOVA, [****] p value<0.0001.

##### Figure S3.

**A.** MYC overexpression vector. **B**. Metabolic flux analysis shows mitochondrial oxygen consumption rates (OCR) reflecting basal mitochondrial oxidative phosphorylation (OXPHOS) in c-MYC-OE or EV transfected MIA PaCa-2 cells following treatment with combination or single drugs. Cells were incubated for a total of 18h including 6h of 15nM romidepsin (both Romi and Romi+DMP) followed by an additional 12h in drug free medium (Romi) or 2nM DMPatA (Romi + DMP) or 18 hours in 2nM DMPatA (DMPatA).

##### Figure S4.

The data in Panels A and B are results of pathway analysis on RNA-seq experiment conducted in MiaPaCa-2 and KLM cells incubated for a total of 18h including 6h of 15nM romidepsin followed by an additional 12h in drug free medium (Romi) or 2nM DMPatA (Romi + DMPatA) or 18 hours in 2nM DMPatA (DMPatA).

Chord diagram showing the linkage between differentially expressed genes after treatment (on the left) and their associated condition (on the right) for a selected pathway. Treatment-related changes in gene expression are indicated in the left outer arc. Number of modified genes per treatment are indicated on the right part of the circle **A**. Change in gene expression in the proton motive force-driven mitochondrial ATP synthesis pathway genes. **B**. Differentially expressed genes in the mitochondrial respiratory chain complex assembly pathway genes.

**Figure S5.**

The data presented in this figure are results of pathway analysis on RNA-seq experiment conducted in MiaPaCa-2 and KLM cells incubated for a total of 18h including 6h of 15nM romidepsin (both Romi and Romi+DMP) followed by an additional 12h in drug free medium (Romi) or in 2nM DMPatA (Romi + DMPatA) or 18 hours in 2nM DMPatA (DMPatA).

**A.** Chord diagram showing the linkage between differentially expressed genes after treatment (on the left) and their associated condition (on the right) for the intrinsic apoptotic signaling pathway genes. Treatment-related changes in gene expression are indicated in the left outer arc. Number of modified genes per treatment are indicated on the right part of the circle.

**Figure S6.**

The data presented in panel A is from pathway analysis on RNA-seq experiment conducted in MiaPaCa-2 and KLM cells incubated for a total of 18h including 6h of 15nM romidepsin (both Romi and Romi+DMP) followed by an additional 12h in drug free medium (Romi) or in 2nM DMPatA (Romi + DMPatA) or 18 hours in 2nM DMPatA (DMPatA).

**A.** Chord diagram showing the linkage between differentially expressed genes after treatment (on the left) and their associated condition (on the right) for the histone modification pathways altered following DMPatA and the combination treatment. Treatment-related changes in gene expression are indicated in the left outer arc. Number of modified genes per treatment are indicated on the right part of the circle. 63 and 183 genes from the histone modification pathways were enriched in the DMPatA and combination treated samples respectively. **B.** Immunoblots showing levels of different HDAC enzymes following treatment of cells with combination (romidepsin 15nM + DMPatA 2nM) and DMPatA as single agent. Romidepsin treatment for all time points was for 6h.

**Figure S7.**

**A.** Immunoblots of H3K27ac (top panel) and H2BK9ac (bottom panel) protein levels from whole cell extracts of MIA PaCa-2 cell line following treatment with combination of romidepsin and DMPatA and single drugs. Romidepsin treatment was for 6h. **B.** Immunoblots of H3K9ac protein levels from whole cell extracts of MIA PaCa-2 and KLM cells untreated or following treatment with combination of entinostat (15μM) and DMPatA (2nM for MIA PaCa-2, and 4nM for KLM cells) and single drugs. Treatments were for various timepoints ranging from 6 to 24h. Entinostat treatment in combination and single drug treatments was only for 6h. **C.** Immunoblots of H3K9ac protein levels from whole cell extracts of MIA PaCa-2 and KLM cells untreated or following treatment with combination of chidamide (15μM) and DMPatA (2nM for MIA PaCa-2, and 4nM for KLM cells) and single drugs. Treatments were for various timepoints ranging from 6- to 24h. chidamide treatment in combination and single drug treatments was only for 6h.

**Figure S8.**

**A**. Immunohistochemical staining of frozen sections from MIA PaCa-2 xenograft tumor tissues using antibodies against H3K9ac, c-MYC, and α-tubulin. The fluoresce signal (red) from H3K9ac and c-MYC antibodies were quantified (see figure 7C-right). DAPI staining shows nuclei of cells. Images are captured at 40x magnification. **B**. Immunoblot of Ɣ-H2AX protein levels from whole cell extracts of wild type and c-MYC over-expressing MIA PaCa-2 cells with or without treatment with single agents or the combination.

Figure S1

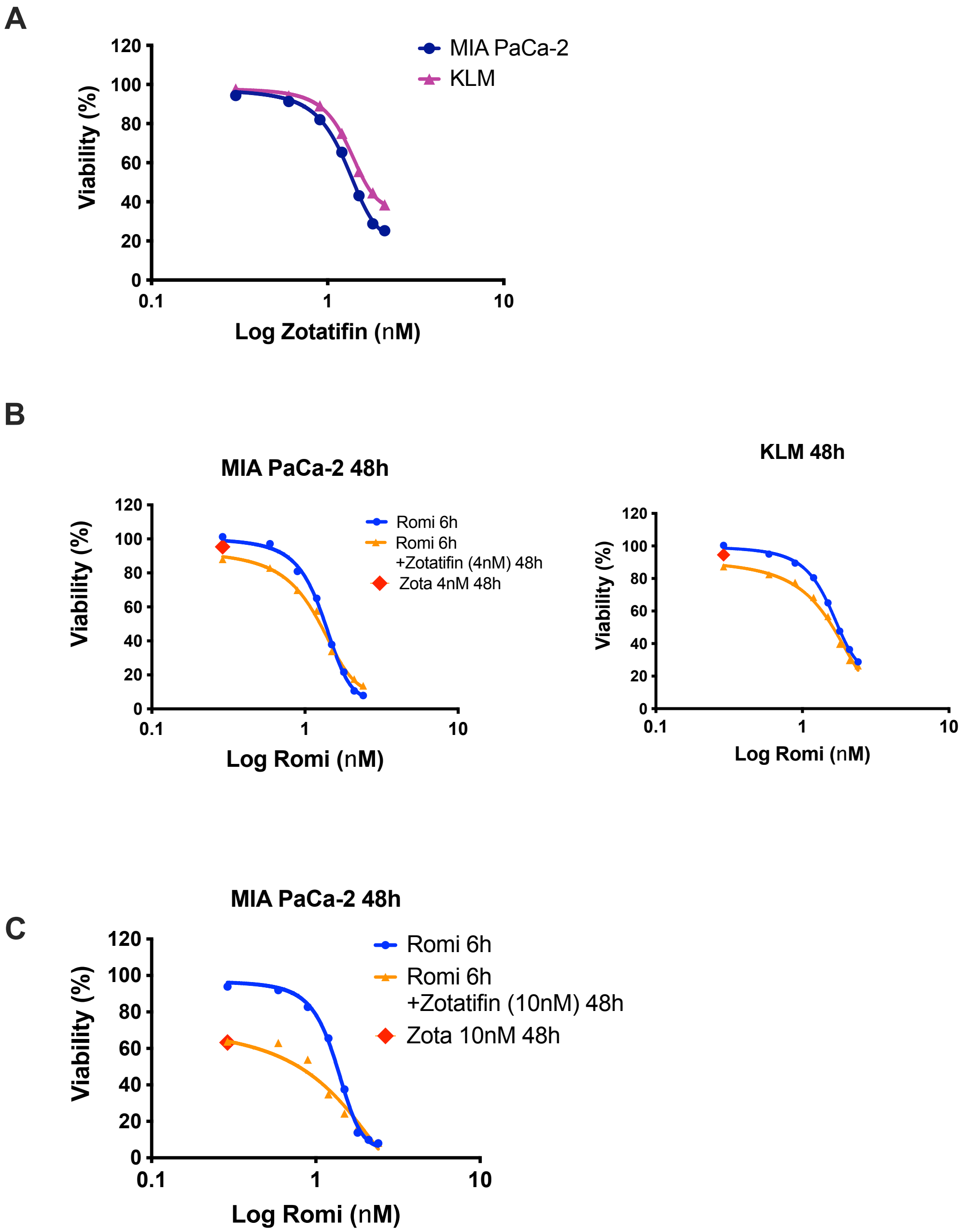
A
MIA PaCa-2
KLM
Viability (%)
Log Zotatifin (nM)
B
MIA PaCa-2 48h
Romi 6h
Romi 6h
+Zotatifin (4nM) 48h
Zota 4nM 48h
Viability (%)
Log Romi (nM)
KLM 48h
Viability (%)
Log Romi (nM)
C
MIA PaCa-2 48h
Romi 6h
Romi 6h
+Zotatifin (10nM) 48h
Zota 10nM 48h
Viability (%)
Log Romi (nM)

Figure S2

A

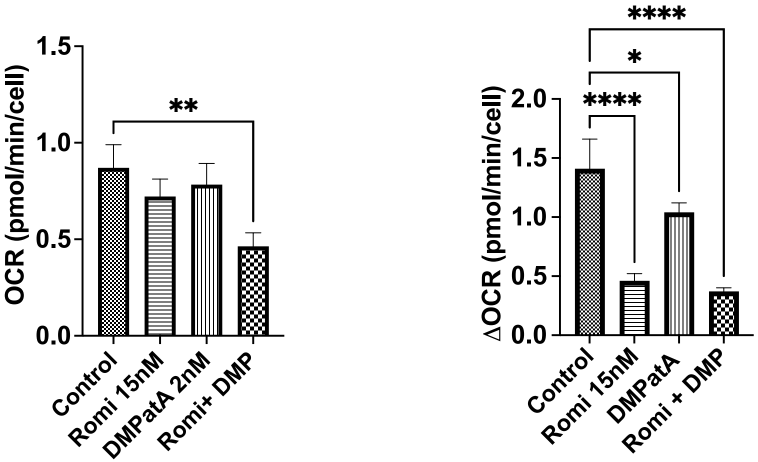
OCR (pmol/min/cell)
1.5
1.0
0.5
0.0
**
Control
Romi 15nM
DMPatA 2nM
Romi+ DMP
ΔOCR (pmol/min/cell)
2.0
1.5
1.0
0.5
0.0
****
*
****
Control
Romi 15nM
DMPatA
Romi + DMP

B

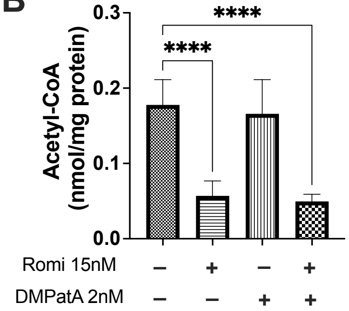
Acetyl-CoA
(nmol/mg protein)
0.3
0.2
0.1
0.0
****
****
Romi 15nM
– + – +
DMPatA 2nM
– – + +

Figure S3

A

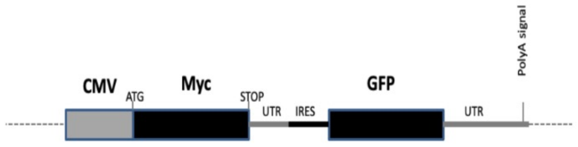
CMV
ATG
Myc
STOP
UTR
IRES
GFP
UTR
PolyA signal

B

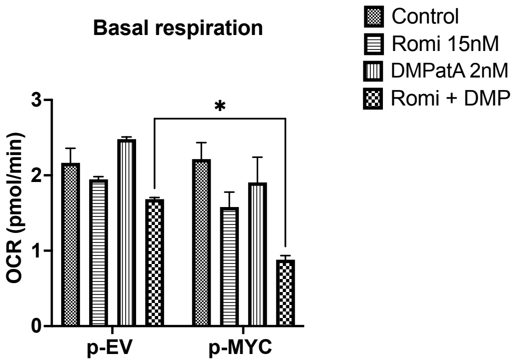
Basal respiration
Control
Romi 15nM
DMPatA 2nM
Romi + DMP
OCR (pmol/min)
3
2
1
0
*
p-EV
p-MYC

Figure S4

**A** Chord diagram _
Proton motive force-driven mitochondrial
ATP synthesis

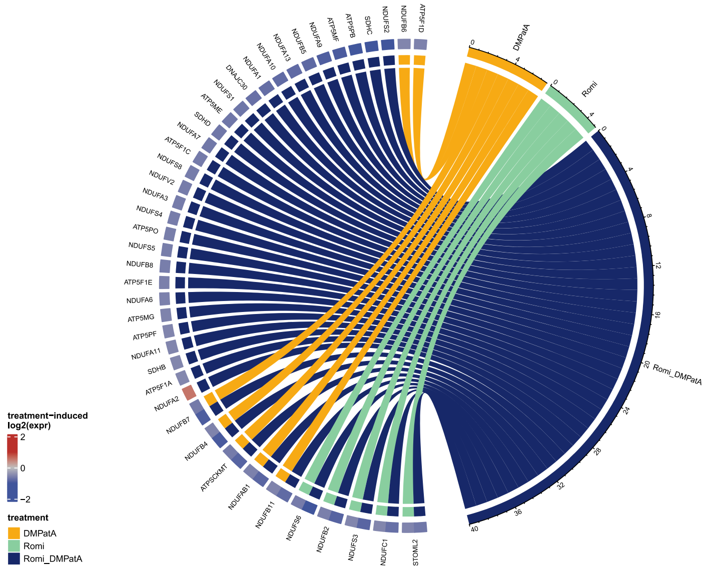


**B** Chord diagram _
Mitochondrial respiratory chain complex assembly

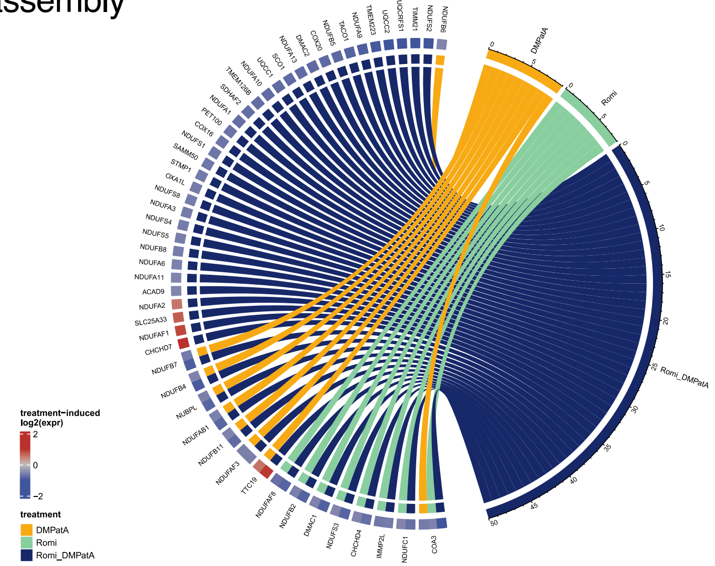

### Figure S5

#### A Chord diagram _Intrinsic apoptotic signaling pathway

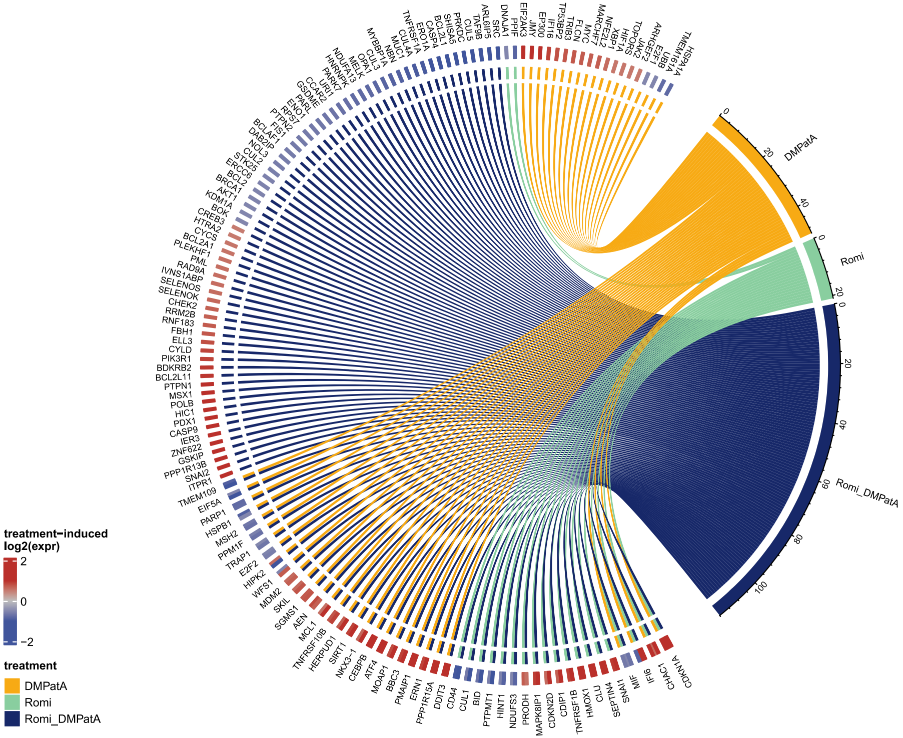

### Figure S6

**A** Chord diagram _Histone modification pathway

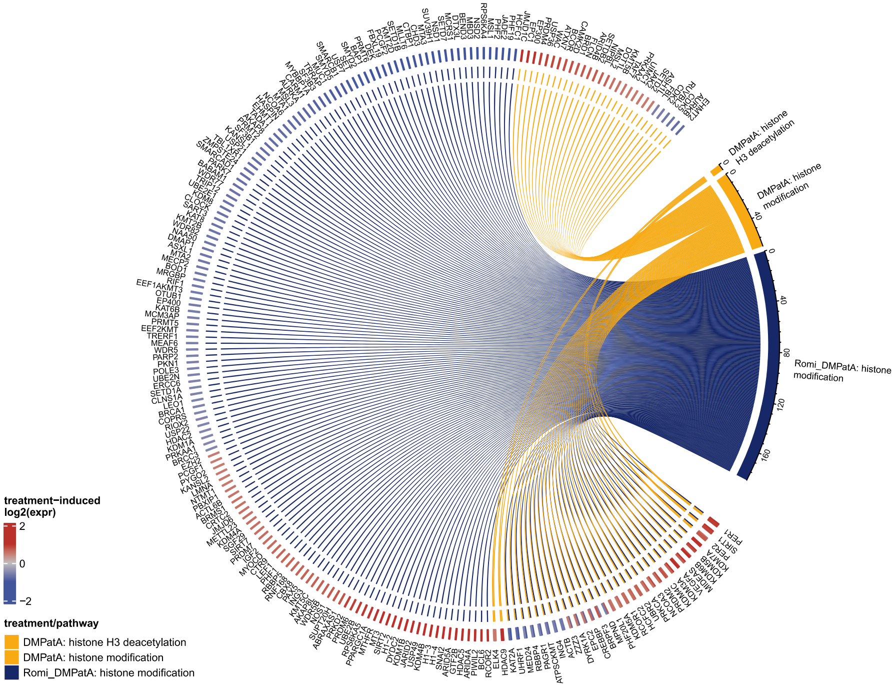


**B**

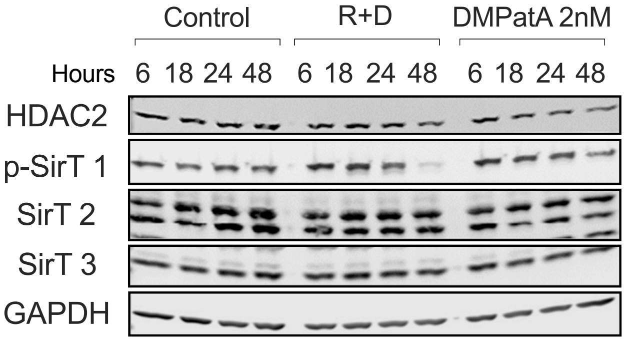

### Figure S7

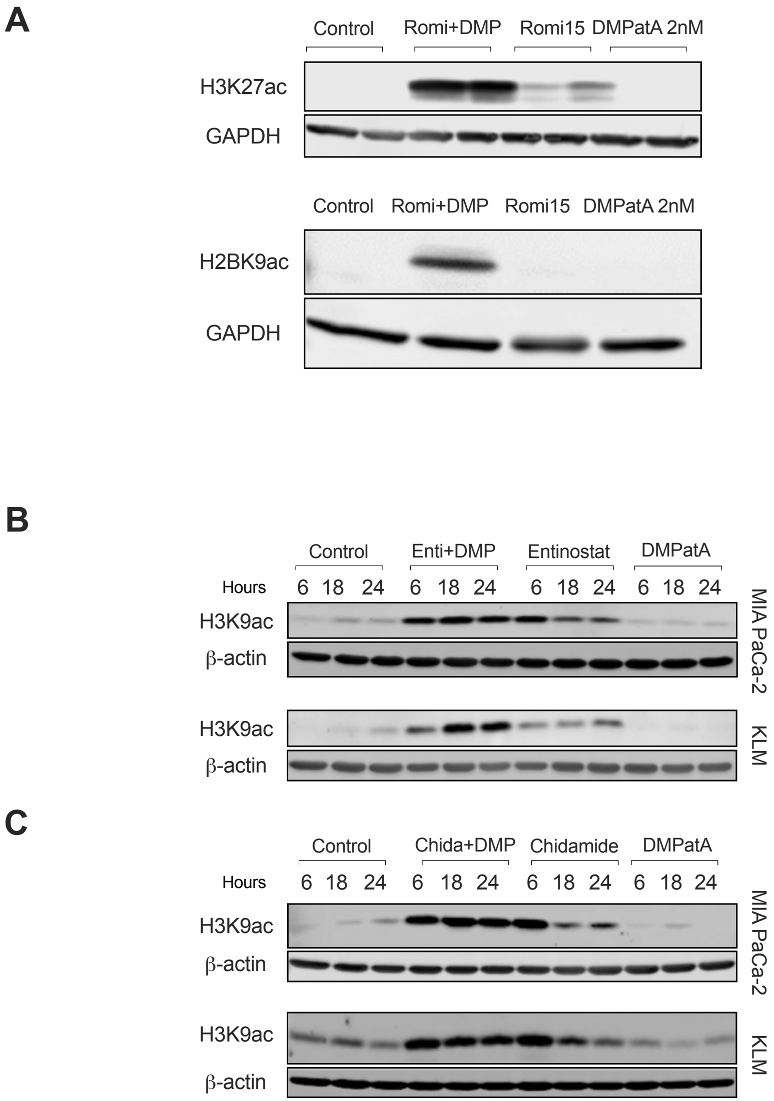
A
Control
Romi+DMP
Romi15
DMPatA 2nM
H3K27ac
GAPDH
Control
Romi+DMP
Romi15
DMPatA 2nM
H2BK9ac
GAPDH
B
Control
Enti+DMP
Entinostat
DMPatA
Hours
6 18 24 6 18 24 6 18 24 6 18 24
H3K9ac
β-actin
MIA PaCa-2
H3K9ac
β-actin
KLM
C
Control
Chida+DMP
Chidamide
DMPatA
Hours
6 18 24 6 18 24 6 18 24 6 18 24
H3K9ac
β-actin
MIA PaCa-2
H3K9ac
β-actin
KLM

### Figure S8

**A**

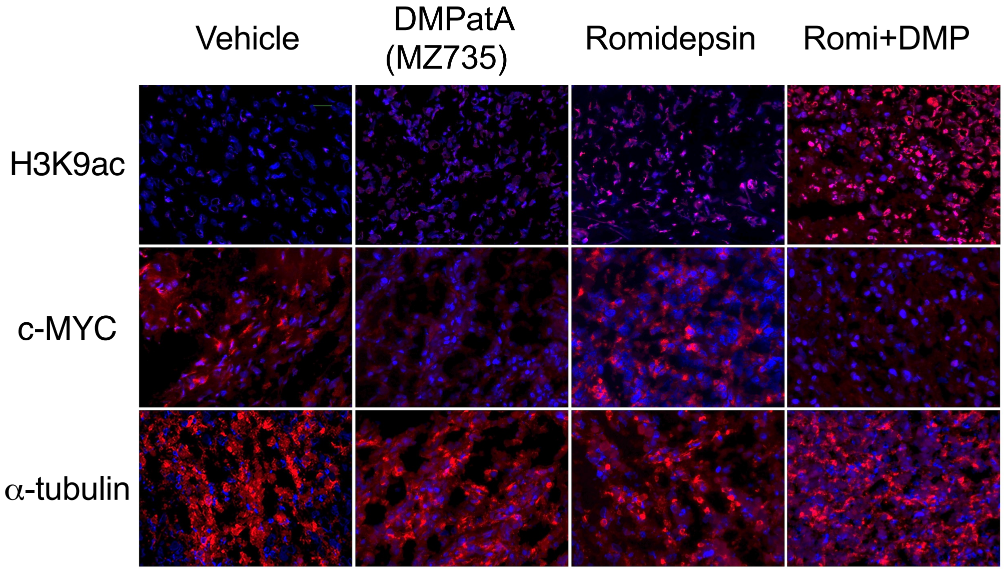

Vehicle
DMPatA (MZ735)
Romidepsin
Romi+DMP
H3K9ac
c-MYC
α-tubulin


**B**

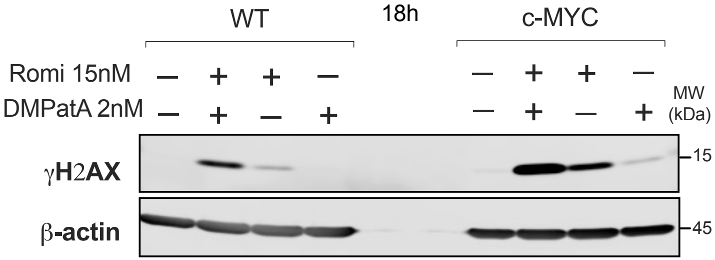

WT
18h
c-MYC
Romi 15nM
DMPatA 2nM
MW (kDa)
γH2AX
β-actin
15
45

**Table S1. Primers used for qPCR validation of DRIP and ChIP**

| Type | Name | Sequence (5'to 3') |
|---|---|---|
| Positive-control locus | *VGF F* | TTCTTCCGCTTCCGCTTC |
| Positive-control locus | *VGF R* | ATTGAGCTGTCCACCAAACT |
| Positive-control locus | *HBA2 F* | ACAGAAGCCAGGAACTTGTC |
| Positive-control locus | *HBA2 R* | CTTCTCTGCACAGCTCCTAAG |
| Negative-control locus | *SNRPN F* | GCCAAATGAGTGAGGATGGT |
| Negative-control locus | *SNRPN R* | TCCTCTCTGCCTGACTCCAT |

### TABLE S2

#### Antibody list

| Name | Company | Cat# | Dilution |
|---|---|---|---|
| Phospho-Histone H2A.X (Ser139) | Millipore Sigma | 05-636 | 1:1000 |
| GAPDH | Abcam | ab8245 | 1:10000 |
| c-Myc | Abcam | ab3207 | 1:1000 |
| β-Actin | Cell signaling Technology | 8457 | 1:1000 |
| Phospho-RPA32/RPA2 (Ser33) | Cell signaling Technology | 10148 | 1:1000 |
| Phospho-Chk1 (Ser345) | Cell signaling Technology | 2341 | 1:1000 |
| HDAC1 (D5C6U) | Cell signaling Technology | 34589 | 1:1000 |
| HDAC2 (D6S5P) | Cell signaling Technology | 57156 | 1:1000 |
| HDAC3 (D2O1K) | Cell signaling Technology | 85057 | 1:1000 |
| SirT1 (D1D7) | Cell signaling Technology | 9475 | 1:1000 |
| Phospho-SirT1 (Ser47) | Cell signaling Technology | 2314 | 1:1000 |
| SirT2 (D4O5O) | Cell signaling Technology | 12650 | 1:1000 |
| SirT3 (D22A3) | Cell signaling Technology | 5490 | 1:1000 |
| IRDye® 680RD Goat anti-Mouse | LI-COR Biosciences | 926-68070 | 1:10000 |
| IRDye® 800CW Goat anti-Rabbit | LI-COR Biosciences | 926-32211 | 1:10000 |
| Mouse anti-BrdU | BD Biosciences | 347580 | 1:25 |
| Rat anti-BrdU | Abcam | ab6326 | 1:400 |
| Goat anti-mouse Cy3 conjugate | Millipore Sigma | AP124C | 1:500 |
| goat anti-rat Alexa Fluor 488 | Invitrogen | A-11006 | 1:400 |
| anti-DNA-RNA hybrid [S9.6] | Kerafast | ENH001 | 1:500 |
| anti-nucleolin antibody | Abcam | ab22758 | 1:1000 |
| Alexa Fluor 488 goat anti-mouse | Invitrogen | A11034 | 1:1000 |
| Alexa Fluor 594 goat anti-rabbit | Invitrogen | A11005 | 1:1000 |
| Normal Rabbit IgG | Cell signaling Technology | 2729 | |
| anti-Acetyl-Histone H3 (Lys9) (C5B11) | Cell signaling Technology | 9649S | 1:1000 (Western blot), 1:50 (ChIP) |
| Anti-phospho-Histone H2A.X (Ser139) | Millipore Sigma | 05-636 | 1:500 |
| Histone H3 (acetyl K9) | Abcam | ab10812 | 1:200 (IHC) |
| C-Myc | Abcam | ab32072 | 1:150 (IHC) |
| alpha tubulin | Abcam | ab7291 | 1:4000 (IHC) |
| Donkey anti-rabbit Alexa 594 | Invitrogen | A21207 | 1:200 |
| Donkey anti-mouse Alexa 594 | Invitrogen | A11032 | 1:200 |
| Acetyl-Histone H3 (Lys27) (D5E4) | Cell signaling Technology | 8173 | 1:1000 |
| Acetyl-Histone H2B (Lys20) (D7O9W) | Cell signaling Technology | 34156 | 1:1000 |
| PARP | Cell signaling Technology | 9542 | 1:1000 |
